# scATAC-Seq reveals epigenetic heterogeneity associated with an EMT-like process in male germline stem cells and its regulation by G9a

**DOI:** 10.1101/2020.10.12.336834

**Authors:** Jinyue Liao, Hoi Ching Suen, Alfred Chun Shui Luk, Annie Wing Tung Lee, Judy Kin Wing Ng, Ting Hei Thomas Chan, Man Yee Cheung, David Yiu Leung Chan, Huayu Qi, Wai Yee Chan, Robin M. Hobbs, Tin-Lap Lee

**Affiliations:** Developmental and Regenerative Biology Program, School of Biomedical Sciences, Faculty of Medicine, The Chinese University of Hong Kong, Shatin, Hong Kong SAR, China; Department of Obstetrics and Gynaecology, Faculty of Medicine, The Chinese University of Hong Kong, Shatin, Hong Kong SAR, China; Guangzhou Regenerative Medicine and Health Guangdong Laboratory, Guangzhou Institutes of Biomedicine and Health, Guangzhou, China; Germline Stem Cell Biology Laboratory, Centre for Reproductive Health, Hudson Institute of Medical Research, Australia

**Keywords:** Single-cell, scATAC-Seq, Epithelial-mesenchymal transition, Spermatogonial stem cell, Chromatin accessibility, G9a, Ehmt2, Male germline

## Abstract

**Background:** Epithelial-mesenchymal transition (EMT) is a phenomenon in which epithelial cells acquire mesenchymal traits. It contributes to organogenesis and tissue homeostasis, as well as stem cell differentiation. Emerging evidence indicates that heterogeneous expression of EMT gene markers presents in sub-populations of germline stem cells (GSCs). However, the functional implications of such heterogeneity are largely elusive.

**Results:** We unravelled an EMT-like process in GSCs by *in vitro* extracellular matrix (ECM) model and single-cell genomics approaches. We found that histone methyltransferase G9a regulated an EMT-like program in GSC *in vitro* and contributed to neonatal germ cell migration *in vivo.* Through modulating ECM, we demonstrated that GSCs exist in interconvertible epithelial-like and mesenchymal-like cell states. GSCs gained higher migratory ability after transition to a mesenchymal-like cell state, which was largely mediated by the TGF-β signaling pathway. Dynamics of epigenetic regulation at the single-cell level was also found to align with the EMT-like process. Chromatin accessibility profiles generated by single-cell sequencing assay for transposase-accessible chromatin (scATAC-seq) clustered GSCs into epithelial-like and mesenchymal-like states, which were associated with differentiation status. The high-resolution data revealed regulators in the EMT-like process, including transcription factors *Zeb1.* We further identified putative enhancer-promoter interactions and *cis*-co-accessibility networks at loci such as *Tgfb1, Notch1* and *Lin28a.* Lastly, we identified HES1 as the putative target underlying G9a’s regulation.

**Conclusion:** Our work provides the foundation for understanding the EMT-like process and a comprehensive resource for future investigation of epigenetic regulatory networks in GSCs.

## Background

Stem cell migration underpins many developmental processes of metazoan animals and germ cell development is no exception. Similar to other stem cells, appropriate migration of spermatogonial stem cells (SSCs) is important for their survival, maintenance and differentiation. Shortly after birth, gonocytes, the precursors of SSCs, resume mitotic activity and begin to migrate to the basal lamina of testicular cords. During a broadly defined timeframe of postnatal days (PND) 0-6, they transition to postnatal spermatogonia, including SSCs[1–3]. In adult testis, SSC migration in seminiferous tubules has also been visualized by intravital live-imaging[4]. Since migrating cells are subjected to different physical microenvironments, spermatogonial migration may be linked to cell fate determination through external signals. A recent study suggested that migrating SSCs could tune their self-renewal and differentiation fate in response to fibroblast growth factor consumption on the basement membrane, suggesting a mechanistic link between SSC differentiation and migration[5]. However, at present, molecular regulatory mechanisms underlying SSC migration as well as its connection with stemness remain largely unknown.

In fact, epithelial-to-mesenchymal transition (EMT) provides an attractive mechanism for explaining how SSCs regulate migration, stemness and the crosstalk between them. EMT is a crucial cellular program that enables polarized epithelial cells to transition toward a mesenchymal phenotype with increased cellular motility. In addition, EMT actively participates in stem cell differentiation during various developmental processes. For example, during gastrulation, epiblast cells undergo EMT to differentiate into three germ layers[6]. During the transition, a hybrid E/M state might exist to allow cells to migrate and respond to external signaling[7]. Classic EMT-transcription factors (EMT-TFs) also participate in developmental programs. Snail regulates neural crest delamination in ectoderm[8].*Zeb2*, together with *Sip1,* modulates the differentiation of embryonic hematopoietic stem cells to progenitor cells[9]. *Zeb2* is also involved in germ layer differentiation from ESCs[10].

Epithelial-mesenchymal (E/M) heterogeneity implies that cells within a population can exhibit epithelial, mesenchymal, or hybrid E/M phenotypes. There is evidence that E/M heterogeneity may exist in undifferentiated spermatogonia, a population of cells containing SSCs and differentiation-primed progenitors. SSCs have generally been considered as neither mesenchymal or epithelial due to expression of both epithelial *(Cdh1)* and mesenchymal *(Thy1)* markers. E-cadherin (CDH1) is a cell-cell adhesion molecule that plays essential roles in epithelial cell behavior and cancer suppression[11], while loss of E-cadherin is a hallmark of EMT[12]. In contrast, THY1, a cell surface glycoprotein, is an adhesion molecule of the immunoglobulin superfamily expressed in fibroblasts and mesenchymal stromal cells[13]. In addition, several SSC-related genes have also implicated roles in EMT processes. For instance, *Cxcr4* and *Foxc2* promote migration and invasiveness through EMT in different types of cancer[14–18]. CD87 (urokinase-type plasminogen activator (uPA) receptor/uPAR), a marker for neonatal undifferentiated spermatogonia, can be regulated by TGFβ and participates in tumor invasion and metastasis[19, 20]. Recent whole transcriptome analysis also confirmed that THY1+ subpopulations of neonatal germ cells were mesenchymal-like due to higher levels of EMT genes *(Vim, Zeb2)* while the ID4+ population (SSC-enriched) expressed lower levels of EMT genes[21].

Whether an EMT or EMT-like process exists in undifferentiated spermatogonia and how it contributes to the regulation of SSC function remain largely unknown. To address this knowledge gap, we searched for potential candidates from our previous single cell RNA-seq study on neonatal undifferentiated germ cells. We found Euchromatic histone-lysine N-methyltransferase 2 (EHMT2), also known as G9a, particularly interesting as it is one of the transcriptional regulators upregulated in differentiation-primed spermatogonia *(Gfra1-/Kit-* cells)[19]. Moreover, neonatal differentiation-primed germ cells expressed higher levels of mesenchymal markers (e.g. *Vim*, *Dab2* and *Mmp3)* along with genes significantly enriched in ECM organization and cell migration[19]. Given the fact that H3K9 methyltransferase G9a is reportedly induced by TGF-β and has an important role in the development of epithelial-to-mesenchymal transition (EMT) in cancer cells[22], we hypothesized that G9a may also play a role in the germ cells through EMT-like mechanisms.

In this study, we characterized an EMT-like process in germline stem cells (GSCs) cultured in vitro, which were derived from PND5.5 Oct4-GFP+/KIT-undifferentiated spermatogonia that contain transplantable SSC populations. First, we found EMT genes were downregulated in GSCs after G9a inhibition. We then recapitulated the EMT-like process on Matrigel through manipulating its concentration in vitro, which revealed the presence of interconvertible epithelial-like and mesenchymal-like GSCs. We confirmed that G9a and the Matrigel culture system shared similar mechanisms in regulating the EMT-like process, as revealed by gene expression and migration capabilities. To account for cellular heterogeneity and revealed transcription factor dynamics in GSC cultures, we leveraged highly multiplexed single-cell ATAC-Seq (scATAC-Seq) to reconstruct the epigenetic regulatory program, reveal distinct cell states during this EMT-like process and identify downstream target of G9a. These data and described strategies provide a useful resource for future epigenetic studies of GSC differentiation and migration.

## Results

### G9a regulates an EMT-like process in germline stem cells

Epithelial/mesenchymal (E/M) heterogeneity has been largely overlooked in current scRNA-Seq studies of spermatogonia, with most studies focused on the regulation of stemness and differentiation. To confirm the heterogeneous expression of G9a, we first examined the published scRNA-Seq datasets which profiled a larger number of cells to reassess the *G9a* expression profile in adult and neonatal spermatogonia[23, 24]. In both sets of data, undifferentiated spermatogonia can be separated into stem cell-like (*Id4*-high) and progenitor cell-like (*Rarg*-high) population, and an additional differentiated-primed cell-like population (*Kit*-high) in neonatal spermatogonia (Supplementary Figure S1A and B). We confirmed that *G9a* mRNA levels were consistently higher in the progenitor-like population compared to stem cell-like population. As the second step to elucidate whether spermatogonia may display E/M or E/M-like heterogeneity, we re-analyzed the EMT-related gene expression in the scRNA-Seq datasets. Interestingly, the progenitor cell-like and differentiation-primed cell-like population showed higher expression of EMT-TFs *(Snail, Zeb1)* and EMT marker genes *(Loxl2, Mmp2)* (Supplementary Figure S1A and B), which confirmed that heterogeneity in EMT-related gene expression exist in spermatogonia.

The loss of cell-cell adhesion molecule CDH1 is considered a hallmark of EMT[12]. To test our hypothesis that G9a affects EMT, we used UNC0638, a substrate-competitive inhibitor that selectively binds enzymatically active G9a, to treat cultured undifferentiated spermatogonia (GSC lines)[25], which were derived from isolated Oct4+/KIT-germ cells of PND5.5 mice and stabilized on mouse embryonic fibroblasts (MEF) as feeder cells (Supplementary Figure S2A and B). We found that UNC0638 treatment for 48 hours resulted in a dose-dependent increase of CDH1+ cells as revealed by FACS analysis (Figure 1A).

**Figure 1.**
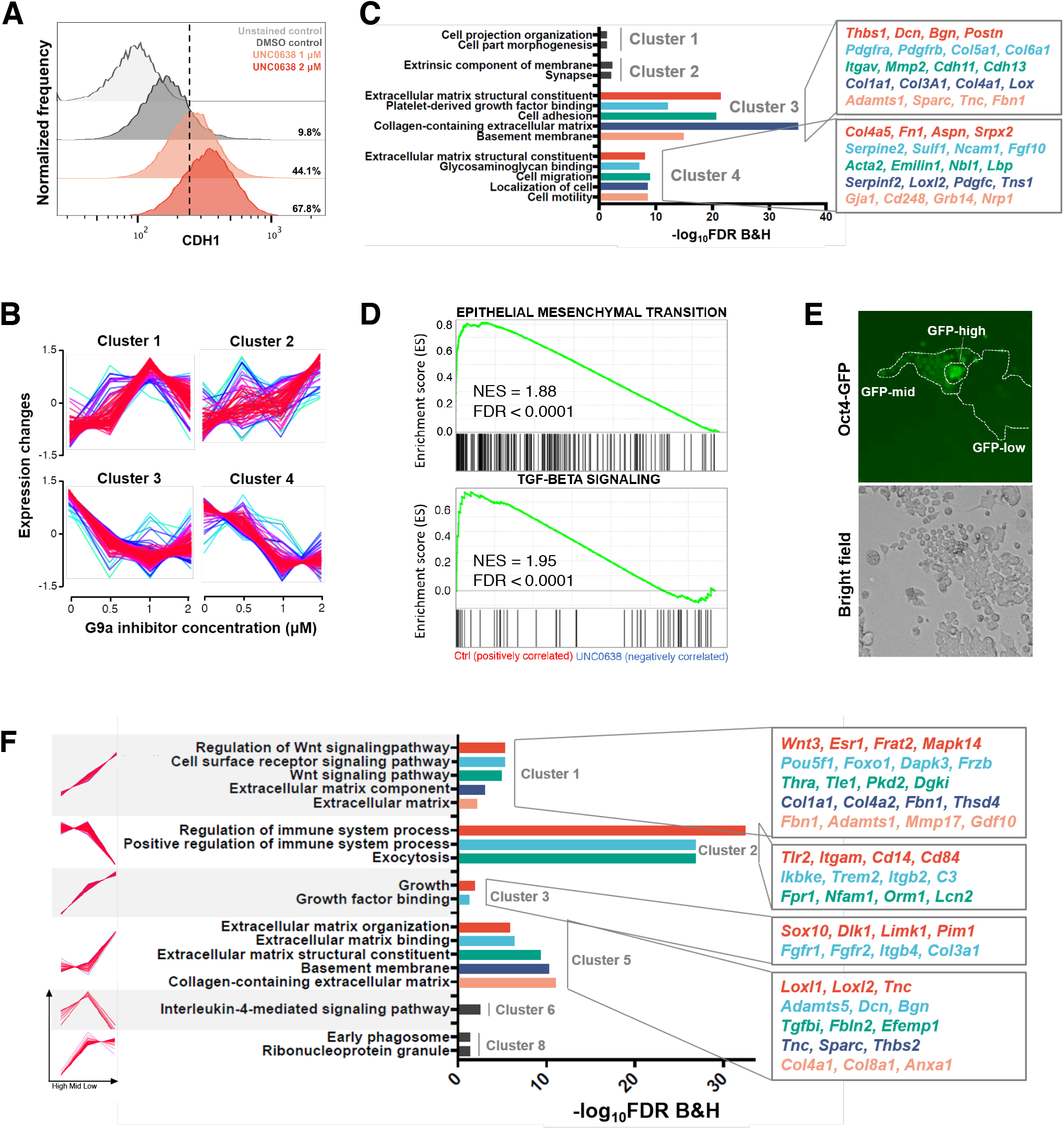
Epithelial-mesenchymal heterogeneity in germline stem cells and its regulation by G9a. **A.** Flow cytometry analysis of epithelial marker CDH1 in UNC0638-treated GSCs. **B.** Differentially expressed genes were identified among the control group and UNC0638 treated groups. Z-score-normalized ratios of the gene expression of 435 DEGs were subjected to soft clustering with Mfuzz software. The 4 clusters represent different expression profiles. **C.** Gene Ontology (GO) terms significantly enriched in each cluster identified from Mfuzz soft clustering (p ≤ 0.05). Representative genes for each cluster are listed in the right column. **D.** GSEA showing the enrichment of EMT Hallmark gene set and TGF-β signaling Hallmark gene set in non-UNC0638-treated GSCs. **E.** Fluorescence images of GSCs cultured on Matrigel show heterogeneity in the Oct4-GFP level. **F.** Gene Ontology (GO) terms significantly enriched in each cluster identified from Mfuzz soft clustering (p ≤ 0.05). Representative genes for each cluster are listed in the right column. Cluster 4 and 7 showed no significant enrichment in GO terms.

To examine the transcriptomic regulation upon G9a inhibition, we treated cells with different dosage of UNC0638 in GSC culture (Supplementary Figure S2C). Mfuzz clustering of differentially expressed genes (DEGs) reveals 4 clusters, in which Cluster 1 (n=79) and Cluster 2 (n=50) showed upregulation while Cluster 3 (n=143) and Cluster 4 (n=61) showed downregulation (Figure 1B, Supplementary Table S1 and S2). Gene Set Enrichment Analysis (GSEA) and GO analysis showed that genes downregulated after UNC0638 treatment were significantly associated with EMT. For example, we observed downregulation of genes related to cell adhesion *(Itgav, Cdh11)* cell motility *(Gja1, Cd248)* and cell migration *(Acta2, Emilin1)* (Figure 1C). We also found several genes related to Wnt signaling pathway and P53 pathway were upregulated after UNC0638 treatment, such as *Axin2, Fzd7, Pten, Siah1, Perp* and *Ep300* (Supplementary Figure S2D). GSEA revealed the most enriched curated gene sets were associated with EMT and TGF-β signaling, which is widely known to induce EMT in many cell types (Figure 1D, Supplementary Figure S2E)[26, 27].

Taken together, these results suggested the possibility that spermatogonia are capable of undergoing an EMT-like process and G9a regulates this process through TGF-β signaling pathway.

### In vitro modelling of epithelial-mesenchymal heterogeneity in neonatal germ cells

Conventional methods to culture germ cells rely on MEF as feeder cells. However, MEF’s ECM properties and its effect on EMT regulation are largely unknown. The presence of a MEF feeder layer also complicates pharmacological manipulation in an in vitro assay. To facilitate the investigation of the functional role of G9a in the EMT-like process, we adopted a Matrigel-based feeder-free culture system as Matrigel can mimic the mechanical and biochemical properties of ECM and support long-term maintenance of GSCs[28]. Cultured GSC lines were first derived and stabilized on MEF, and then transferred to Matrigel-coated plates for subsequent long-term culture. To address whether the cultured cells still maintain a self-renew gene expression program, cells were examined during 9 days of the MEF-to-Matrigel transition period (Supplementary Figure S3A). We confirmed GSCs in Matrigel-based culture expressed comparable levels of a panel of stem *(Gfra1, Etv5)* and progenitor-associated genes *(Upp1, Rarg)* as MEF-based cultures throughout the transition period (Supplementary Figure S3B).

To examine whether E/M-like heterogeneity observed in vivo is maintained in the culture system [19], we separated GSC subpopulations from culture based on Oct4-GFP intensity by FACS and performed RNA-Seq (Figure 1E, Supplementary Figure S3C). Consistent with expression patterns in vivo from our previous study, stemness gene expression was higher in the Oct4-GFP-low subpopulation, while progenitor gene expression was higher in the Oct4-GFP-high subpopulation (Supplementary Figure S3D) [19]. These observations are also consistent with a previous study that analysed spermatogonial cultures derived from adult Oct4-GFP mice [23]. We observed that the Oct4-GFP-high subpopulation exhibited upregulation of *Tgfb1, Ehmt2* and well-known mesenchymal markers, such as *Mmp2* and *Loxl2*, although there was no significant difference in EMT-TF expression (*Zeb*, *Snail)*(Supplementary Figure S3E). Of the 589 differentially expressed genes (DEGs) (FDR < 0.05 and fold change ≥ 2) identified, 371 DEGs were upregulated in Oct4-GFP-high GSCs while 218 were upregulated in Oct4-GFP-low GSCs (Supplementary Table S3). Mfuzz clustering analysis reveals 8 clusters of different expression trends (Supplementary Table S4, Supplementary Figure S3F). The GO terms enriched in DEGs in Cluster 1 (n=103) and Cluster 5 (n=49) with positive correlation to Oct4-GFP expression were associated with Wnt signaling and ECM (Figure 1F). For example, Cluster 1 DEGs include metalloprotease genes

*Adamtsl* and *Mmp17,* which are known to participate in cancer progression[29, 30]. Cluster 3 showed a similar trend and enriched GO terms were related to growth factor binding. Among them, FGFR2 mediates mouse SSC self-renewal through MAP2K1 signaling[31]. On the other hand, the DEGs in Cluster 2 (n=145) with negative correlation to Oct4-GFP levels are enriched for immunity. Collectively, the bulk RNA-seq data indicated Oct4-GFP-high cells might contain a subpopulation of cells with a more mesenchymal-like gene expression program in both in vitro and in vivo condition. Our culture system thus successfully recapitulated E/M-like heterogeneity of germ cells.

### Modulation of epithelial-mesenchymal plasticity of GSCs through controlling ECM properties

Despite the association between high Oct4-GFP level and a more mesenchymal-like property, we could not neglect the fact that high Oct4-GFP level is observed in both gonocytes (precursors of SSC) and differentiating spermatogonia. It is therefore not desirable to investigate the E/M-like heterogeneity in germ cells solely based on Oct4-GFP level. The aforementioned upregulation of genes related to ECM and ECM organization may lead to increase in ECM components and promote EMT induction [32]. To test this possibility in spermatogonia and establish an effective in vitro model, we modulated ECM components using different concentrations of Matrigel coating. Strikingly, we observed the formation of two morphologically distinct GSC colonies with similar cell proliferation rates: low (0.1-1 μg/cm^2^) and high (5-20 μg/cm^2^) concentrations of Matrigel promoted the formation of domed-shaped colonies and flat-shaped colonies respectively (Figure 2A, Supplementary Figure S4A). Domed GSCs displayed a more epithelial-like phenotype and were packed into a round colony, possibly due to higher cell-cell adhesion. In contrast, flat GSCs exhibited a more mesenchymal-like phenotype, marked by more cell protrusions. RT-PCR analysis indicated that GSCs cultured on 0.1 μg/cm^2^ of Matrigel expressed higher levels of epithelial marker *Cdh1* (encoding E-cadherin), while GSCs cultured on 20 μg/cm^2^ of Matrigel expressed higher levels of the mesenchymal marker *Vim* (encoding vimentin) (Supplementary Figure S4B).

**Figure 2.**
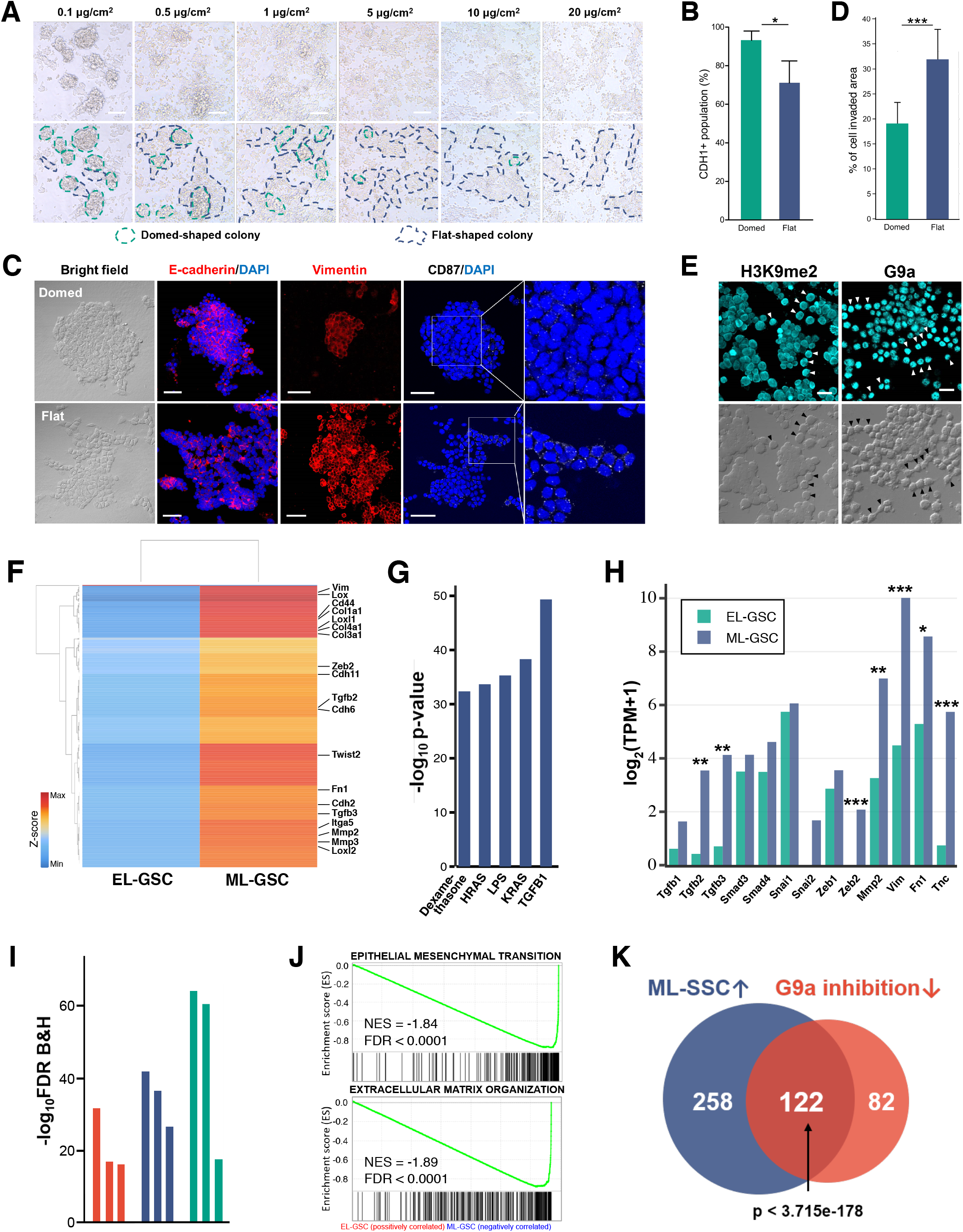
Gene expression signatures of epithelial-like (EL) and mesenchymal-like (ML) germline stem cells (GSCs). **A.** Bright-field images of GSC cultured on different concentrations of Matrigel coating after 14 days showing distinct domed and flat morphology. Scale bars: 100 μm. **B.** Quantification of CDH1+ proportion in domed GSCs and flat GSCs by flow cytometry analysis (p < 0.05, n = 3). Significance was determined by an unpaired two-tailed t-test. **C.** Representative bright-field and immunofluorescence images of domed and flat GSC colonies from at least 5 images per group. Scale bars: 20 μm. **D.** Matrigel-precoated membrane filter insert was used to measure in vitro invasiveness. After 48 hours of incubation, the area invaded by cells was measured. Quantification of invasive cells (p < 0.001, n ≥ 3). Significance was determined by an unpaired two-tailed t-test. **E.** Representative bright-field and immunofluorescence images of Matrigel-based GSC culture stained for G9a and H3K9me2 (from at least 5 images per group). The arrowheads indicate flat cells with stronger G9a and H3K9me2 protein levels. Scale bars: 20 μm. **F.** Heatmap of 381 differentially expressed genes (DEGs) identified by comparing EL-GSC and ML-GSC (p < 0.05, FC > 2). **G.** Top 5 upstream regulators predicted by Ingenuity pathway analysis (IPA). **H.** The expression level of genes involved in TGFβ-induced EMT pathway in EL-GSCs and ML-GSCs. * FDR < 0.01. ** FDR < 0.05. *** FDR < 0.001. **I.** Gene ontology (GO) terms significantly enriched for DEGs upregulated in ML-GSCs compared with EL-GSCs (Red: Molecular Function, Blue: Biological Process, Green: Cellular Component) (p ≤ 0.05). **J.** GSEA showing the enrichment of EMT Hallmark gene set and Extracellular matrix organization gene set in ML-GSC compared to EL-GSC. **K.** Venn diagram showing significant overlapping of DEGs that upregulated in ML-GSC (ML-GSC vs EL-GSC) and downregulated after G9a inhibition.

Domed GSC colonies were less firmly attached to the matrigel and would detach after gentle pipetting, which allowed easy separation of domed and flat GSCs for further characterization. Flow cytometry analysis indicated that the CDH1+ proportion of domed GSCs is 93.2%, which is significantly higher than that of flat GSCs (71.1%) (n=3, p<0.05), while the CDH1+ proportion of GSC on MEF was at an intermediate level (82.1%) (Figure 2B, Supplementary Figure S4C). Consistent with this, immunofluorescence staining revealed that flat GSCs expressed higher levels of vimentin, while domed GSCs showed higher E-cadherin levels (Figure 2C). Interestingly, we found that flat colonies expressed higher levels of CD87 (uPAR), which has been suggested to be involved in germ cell migration (Figure 2C)[19]. Indeed, transwell invasion assay showed that flat GSCs had higher invasive capacity (Figure 2D, Supplementary Figure S4D). Based on their characteristics, we applied the term epithelial-like GSCs (EL-GSC) to refer to domed GSCs and the term mesenchymal-like GSCs (ML-GSC) to refer to flat SSCs.

We further examined the levels of G9a protein and G9a-mediated H3K9me2 modification in our in vitro Matrigel-based GSC culture. The levels of G9a and H3K9me2 in ML-GSCs (black arrows, cells with more protrusions) was higher than that of EL-GSCs, which supported our previous results showing G9a promoted the EMT-like process in GSCs (Figure 2E).

To assess the ability of GSCs to transition between the epithelial-like and mesenchymal-like states, we cultured flat GSCs on low concentration Matrigel and monitored them for formation of domed colonies. Following re-plating of cell suspensions, domed-shaped colonies became apparent over a period of 14 days. Likewise, cells derived from domed colonies on low matrigel concentrations formed flat colonies on high concentration Matrigel over this time frame (Supplementary Figure S4E).

Taken together, these results demonstrated that GSCs could undergo EMT- or MET-like processes in vitro through modulating ECM property.

### Shared mechanisms of the activation of EMT-like program and G9a regulation in GSC

To investigate the underlying regulatory pathways during GSC EMT/MET-like process, we performed bulk RNA-Seq analysis on EL-GSCs and ML-GSCs and identified 381 DEGs with FDR < 0.05 and fold change ≥ 2. Surprisingly, most of the DEGs were upregulated in ML-GSCs (Figure 2F, Supplementary Table S5). Of these, many were key players of the EMT pathway. We then examined the top upstream regulators in ML-GSCs predicted by Ingenuity Pathway Analysis (IPA) (Figure 2G). TGFB1 caught our attention as many of its interacting partners were also upregulated in ML-GSC, such as CDH2, LOXL2 and MMP2, and G9a caused upregulation in TGF-β signaling genes (Figure 1D, Supplementary Figure S5A). We also took a closer look at genes that are involved in TGF-β-induced EMT pathway, such as *Tgfb1*, *Zeb2* and *Snai2*. In accordance with IPA results, we found that most of the genes involved in TGF-β-induced EMT were also upregulated in ML-GSCs (FDR < 0.05) (Figure 2H). We then used IPA to determine the top significant networks upregulated in ML-GSC. The key nodes in the highest scored network are collagens and ERK1 (Supplementary Figure S5B). The activation of ERK1/2 pathway promotes TGFβ-induced EMT and cancer metastasis[33]. The network with second highest score infers NF-kB pathway, which is another pathway essential for both the induction and maintenance of EMT (Supplementary Figure S5C) [34].

The GO terms significantly enriched in ML-GSC included “collagen-containing extracellular matrix”, “extracellular structure organization” and “biological adhesion” (Figure 2I). Genes enriched in GO term “biological adhesion”, including *Fn1* (encoding fibronectin) and *Itga5*(encoding integrin β4), have also been implicated in cell invasion and migration during EMT [35, 36]. GSEA validated that the most enriched curated gene sets were associated with ECM proteins and regulators, such as “Core Matrisome”and “Extracellular Matrix Organization” (Figure 2J, Supplementary Figure S5D). Furthermore, the EMT hallmark gene set was enriched in ML-GSC, indicating ML-GSCs were potentially GSCs that had undergone an EMT-like transition (Figure 2J).

We further validated the link between G9a and EMT in GSC. UNC0638-inhibited DEGs significantly overlapped with the DEGs upregulated in ML-GSC (p < 3.715e-178, Fisher’s exact test), consisting of a prominent set of EMT genes associated with the activation of TGF-β signaling such as *Vim*, *Fn1* and *Twist2* (Figure 2K, Supplementary Figure S6A). GSEA verified that the upregulated DEGs in ML-GSC were significantly enriched in the control group when compared to G9a treatment group (p < 0.0001) (Supplementary Figure S6B). Since cells that undergo EMT often acquire increased migratory capabilities, we questioned whether this also applied to spermatogonia. Thus, we further examined the gene set preferentially expressed in the recently discovered migratory pro-spermatogonia (I-ProSG), which presumably regulates gonocyte migration in vivo[37]. GSEA analysis revealed enrichment of this gene set in ML-GSC and UNC0638-untreated cells respectively, with core-enriched genes related to cell adhesion (Supplementary Figure S6C to E), implying the association among the role of G9a, EMT-like process and migration abilities in spermatogonia.

### The EMT-like process promotes cell migration in spermatogonia and its regulation by G9a

To directly assess the impact of G9a on EMT-like progression and cell migration, we devised an in vitro assay in which cells from a domed-shaped colony (EL-GSC) migrate into unoccupied margins of the well (Figure 3A). We seeded GSCs on low concentration Matrigel to induce domed-shaped colonies. We then transferred the colonies to high concentration Matrigel after which cells at the colony border could spontaneously undergo EMT-like process and migrate outwards. Cells at the periphery of the patch acquired a spindle-like morphology and formed leading and protruding edges consistent with the acquisition of a mesenchymal phenotype (Supplementary Figure S6F). To demonstrate the robustness of this assay, we examined the effect of several kinase inhibitors. Consistent with the upregulation of the TGF-β pathway in ML-GSC, spontaneous EMT-like progression depends on TGF-β, as TGF-β type 1 receptor inhibitor RepSox could significantly inhibit outward migration (Figure 3B). We also validated the action of other potential kinases *(Pdgfrb, Fgfr1, Flt4, Egfr),* whose gene expression was upregulated in ML-GSCs. These kinases are also reported to have roles in EMT and cancer metastasis[38–41]. Indeed, their respective inhibitors suppressed GSC migration (Supplementary Figure S6G and H). More importantly, we observed similar suppression of EMT-like progression when domed-shaped EL-GSC colonies were treated with G9a inhibitor UNC0638 (Figure 3B).

**Figure 3.**
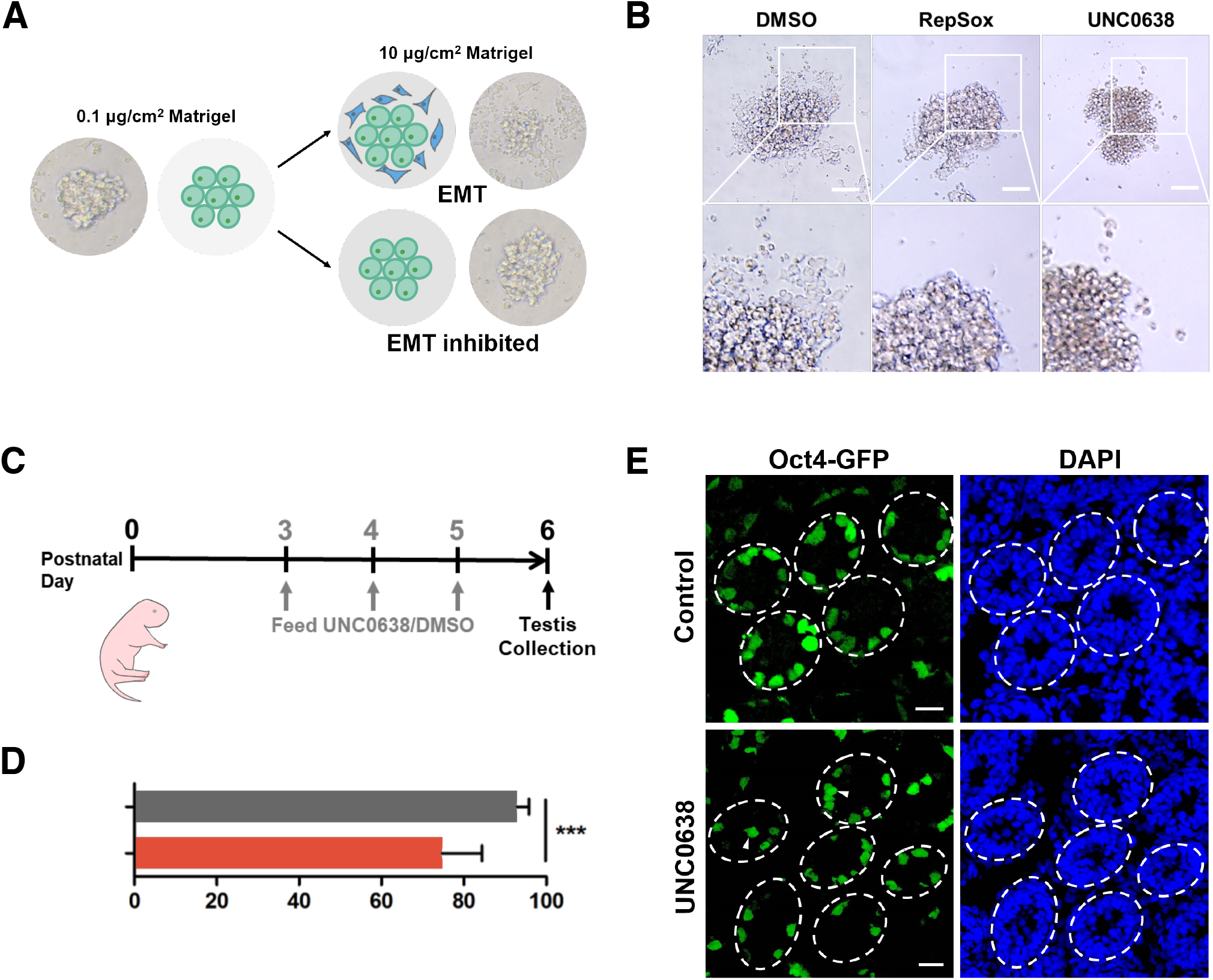
The EMT-like process promotes cell migration in spermatogonia. **A.** A schematic diagram of in vitro assay for studying spontaneous EMT progression. Domed GSC colonies (green) are isolated from low concentration Matrigel culture and then transferred to high concentration Matrigel culture. The colonies are subsequently treated with small molecule inhibitors and cell migration is monitored (blue). **B.** Bright-field images of domed GSC colonies treated with small molecule inhibitors (RepSox 5 μM, UNC0638 2 μM) or vehicle (DMSO) for 24 hours. Dash lines indicate colony boundary at 0 hour. **C.** Schematic diagram of in vivo G9a inhibition. UNC0638 was administered in neonatal mice on PND3-5 and samples were collected on PND6. **D.** Representative confocal images of testis sections from UNC0638-treated and control (DMSO) mice at PND6. Oct4-GFP expression was observed in undifferentiated germ cells. Cell nuclei were stained with DAPI. Arrows indicate germ cells not attached to the basement membrane. Scale bars: 20 μm. **E.** Quantification of percentage of Oct4-GFP+ spermatogonia attached to basement membrane on PND6 (p < 0.001, UNC0638 = 6 testes, DMSO = 3 testes). Significance was determined by an unpaired two-tailed t-test.

Next, we studied the effect of G9a to germ cell migration in vivo. We administered UNC0638 daily by oral gavage (p.o.) to neonatal male mice from PND3 for three constitutive days and observed its effect (Figure 3C). On PND6, the testes were collected and proportion of Oct4-GFP+ nascent spermatogonia attached to the basement membrane were quantified. Most of the Oct4-GFP+ cells (92.9%, n=3) finished migration on PND6 in the control group.

However, in UNC0638-treatment group, significantly fewer Oct4-GFP+ (74.7%, n=6, p<0.001) cells attached to the basement membrane and a large portion of cells remained at the centre of the tubule lumen (Figure 3D and E). This demonstrated that G9a is required for proper migration of GSC.

Collectively, both in vitro and in vivo evidence suggested that G9a might be involved in germ cell migration through EMT-related mechanisms.

### Single cell chromatin states define EMT spectrum in GSC

EMT is dynamically orchestrated by transcription factors (TFs) in both developmental and tumor settings[42]. ATAC-Seq can capture the chromatin accessibility and allows elucidation of TF activity. However, bulk ATAC-Seq may miss important biological signals carried by only a subset of cells in heterogeneous populations of GSC. Therefore, we used scATAC-Seq to investigate both cellular heterogeneity and dynamic TF regulation during the GSC EMT/MET-like transition. We performed scATAC-Seq of the followings: 1) GSCs treated with UNC0638; 2) GSCs treated with vehicle (DMSO); 3) Matrigel-based GSC culture undergoing “forced” EMT-like process containing a pool of cells collected from 3 time points (1, 7 and 14 days) after they were transferred to high concentration Matrigel coating (Figure 4A). After filtering out low-quality nuclei and doublets, we obtained a total of 9,969 high-quality single nuclei for downstream analysis (Supplementary Figure S7A).

**Figure 4.**
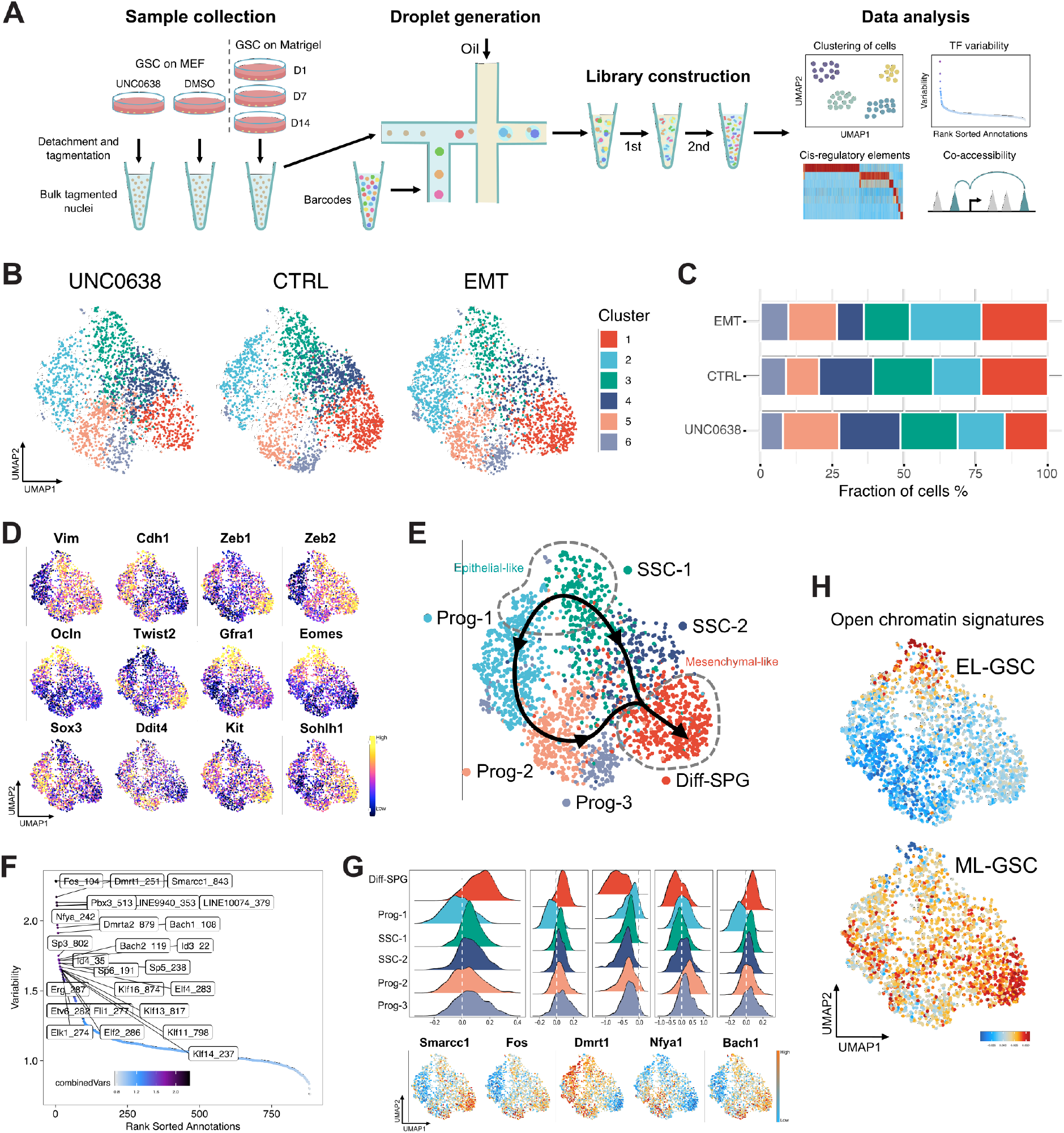
Single-cell chromatin state of germline stem cells (GSCs). **A.** The workflow of scATAC-seq. Samples were first collected from culture and subjected to tagmentation. The libraries were then constructed on on BioRad SureCell ATAC-Seq platform to measure single nuclei accessibility, followed by downstream data analysis by ArchR package. **B.** Uniform manifold approximation and projection (UMAP) projection of scATAC-seq profiles of 1) GSCs treated with UNC0638 (UNC0638); 2) GSCs treated with vehicle (CTRL); 3) Matrigel-based GSC culture containing a pool of cells collected from 3 time points (EMT), generated by SnapATAC. **C.** Bar chart showing the distribution of cells in each cluster. **D.** Gene activity score of EMT marker genes and SSC self-renew and differentiation genes shown in UMAP. **E.** De novo identification of cell states visualized using UMAP. **F.** Variability of TF motifs calculated by chromVAR. **G.** Violin plot (top) and UMAP (bottom) showing the motif accessibility of five of the most variable motifs. **H.** Open chromatin signatures of EL-GSC and ML-GSC identified from bulk ATAC-seq were enriched in distinct locations shown in UMAP.

We adopted SnapATAC to cluster the cells based on the similarity of the chromatin accessibility landscape, which is one of the preferred computational pipelines in recent benchmarking studies[43, 44]. SnapATAC also clusters cells using the cell-by-bin matrix which makes it easy to combine multiple samples and perform comparative analysis. SnapATAC portioned these nuclei profiles into six clusters and the clustering results were not affected by sequencing depth (Figure 4B, Supplementary Figure S7B). The three samples showed overall similar distributions of the six clusters with varied proportions (Figure 4C).

We reasoned that the “forced” EMT-like condition would capture cells from different stages during transition and offer a more holistic view of the whole EMT-like process. Thus we first defined the differentiation state and EMT spectrum in GSC by downstream analysis on this sample. First, we exploited gene scores to explore cellular properties. A gene score is essentially a prediction of how highly expressed a gene will be based on the accessibility of regulatory elements in the vicinity of the gene, including the gene body and putative distal regulatory elements. SSC-1 is likely to be SSC enriched, as most cells in this subset expressed previously recognized mouse SSC marker genes, including *Gfra1* and *Eomes* (Figure 4D). Cells in SSC-2 also expressed *Gfra1* and *Eomes* but at a lower level. Prog-1 cells selectively express progenitor markers *Sox3, Nanos3* and *Neurog3* (Figure 4D, Supplementary Figure S7C). Prog-2 and Prog-3 share a similar gene expression profile, with the expression of progenitor genes *Ddit4.* Lastly, we found increased gene scores of differentiation markers *Kit* and *Sohlh1* in the Diff-SPG subset (Figure 4D). We then explored the existence of an EMT-like spectrum among the 6 spermatogonia subsets. A subset of cells from SSC-1 and Prog-1 located at the upper region of the UMAP embedding shared higher levels of epithelial markers *Cdh1* and *Ocln,* which we referred to as epithelial-like state (EL-state). Consistently, this subset of cells displayed lowest levels of mesenchymal markers *Vim* and EMT-TFs such as *Zeb1, Zeb2, Snail* and *Snai2* (Figure 4D, Supplementary Figure S7D). In contrast, Diff-SPG subset was predicted to express lower levels of epithelial markers *Cdh1* and *Ocln* and increased levels of EMT-TFs (Figure 4D), which we referred to as mesenchymal-like state (ML-state). Thus, cell clustering based on chromatin accessibility revealed cell states spanning GSC differentiation and EMT-like transition (Figure 4E).

Second, we inferred TF motif activity (or in short, TF activity) from scATAC-Seq data. ChromVAR calculates a TF z-score, which infers the binding of TF to open chromatin regions based on the presence of the TF motifs within these regions[45]. We found that the TF z-score of core EMT-TFs *Zeb1, Twist2, Snai1* and *Snai2* are higher in Diff-SPG, supporting that Diff-SPG are more mesenchymal-like (Supplementary Figure S7E). In addition, chromVAR analysis showed that the most variable TF motifs across all the cells represented EMT regulators *Smarcc1, Bach1* and *c-Fos* (Figure 4F and G). BACH1 has been reported to promote EMT by repressing the expression of a set of epithelial genes [46]. SMARCC1 interacts with NKX6.1 to activate EMT in HeLa cells[47]. FOS forms a multimeric SMAD3/SMAD4/JUN/FOS complex to regulate TGF-β-mediated signaling upon TGF-β activation[48]. Notably, these factors display higher activity in Diff-SPG, suggesting their possible role at the later stage of EMT. In addition, a decreased DMRT1 TF z-score was found in Diff-SPG, which is consistent with the previous finding that loss of DMRT1 promoted spermatogonial differentiation through increasing RA responsiveness and activating *Sohlh1* transcription[49].

Third, we tried to associate the chromatin accessibility and phenotypic information. We reasoned that chromatin accessibility signatures associated with bulk populations of EL- and ML-GSCs should preferentially be enriched at either end of the EMT-like spectrum in single cell data. To test this, we performed bulk ATAC-Seq on EL- and ML-GSCs. We first confirmed that EL-GSCs showed higher chromatin accessibility in epithelial marker genes and lower chromatin accessibility in mesenchymal markers and EMT regulators, compared to ML-GSCs (Supplementary Figure S8). Using an analytical framework similar to chromVAR to evaluate enrichment of differentially accessible regions (DARs) between these two phenotypes, we found that EL-GSC and ML-GSC signatures were clearly enriched in ELand ML-state, respectively (Figure 4H).

Taken together, single-cell chromatin accessibility profiling successfully deconvoluted cellular epigenetic states associated with both differentiation and EMT-like process in GSC. Our results suggested that chromatin accessibility dynamics associated with GSC differentiation are coupled with E/M heterogeneity: stem cells are more epithelial-like while differentiating cells are more mesenchymal-like, which is reminiscent of the finding from in vivo scRNA-seq as described above.

### Dynamic changes in accessibility of regulatory elements and TF activity during the EMT-like process

While scRNA-seq has revealed the continuous nature of spermatogonial differentiation through transcriptome analysis, we hypothesized that chromatin accessibility could highlight an additional and distinct layer of transcription factor dynamics. We adopted a semi-supervised pseudotime approach to reconstruct the EMT-like trajectory of GSCs. We reasoned that stem cells at the start of the trajectory would transverse through intermediate states before generating differentiating cells. Given the close association between GSC differentiation and EMT-like processes, this path should also chart a trajectory from the EL- to ML-state. 3D UMAP embedding suggested two possible trajectories as cell fates appeared to diverge via two distinct branches from SSC-1 (Figure 5A, Supplementary Figure S7F). The state sequence in Trajectory 1 corresponds to SSC-1 (EL-state) differentiating into progenitors (Prog-1/2/3) and later into Diff-SPG (ML-state), while in Trajectory 2, SSC-1/2 bypassed the intermediate progenitor-state and directly reached the differentiating state (Diff-SPG).

**Figure 5.**
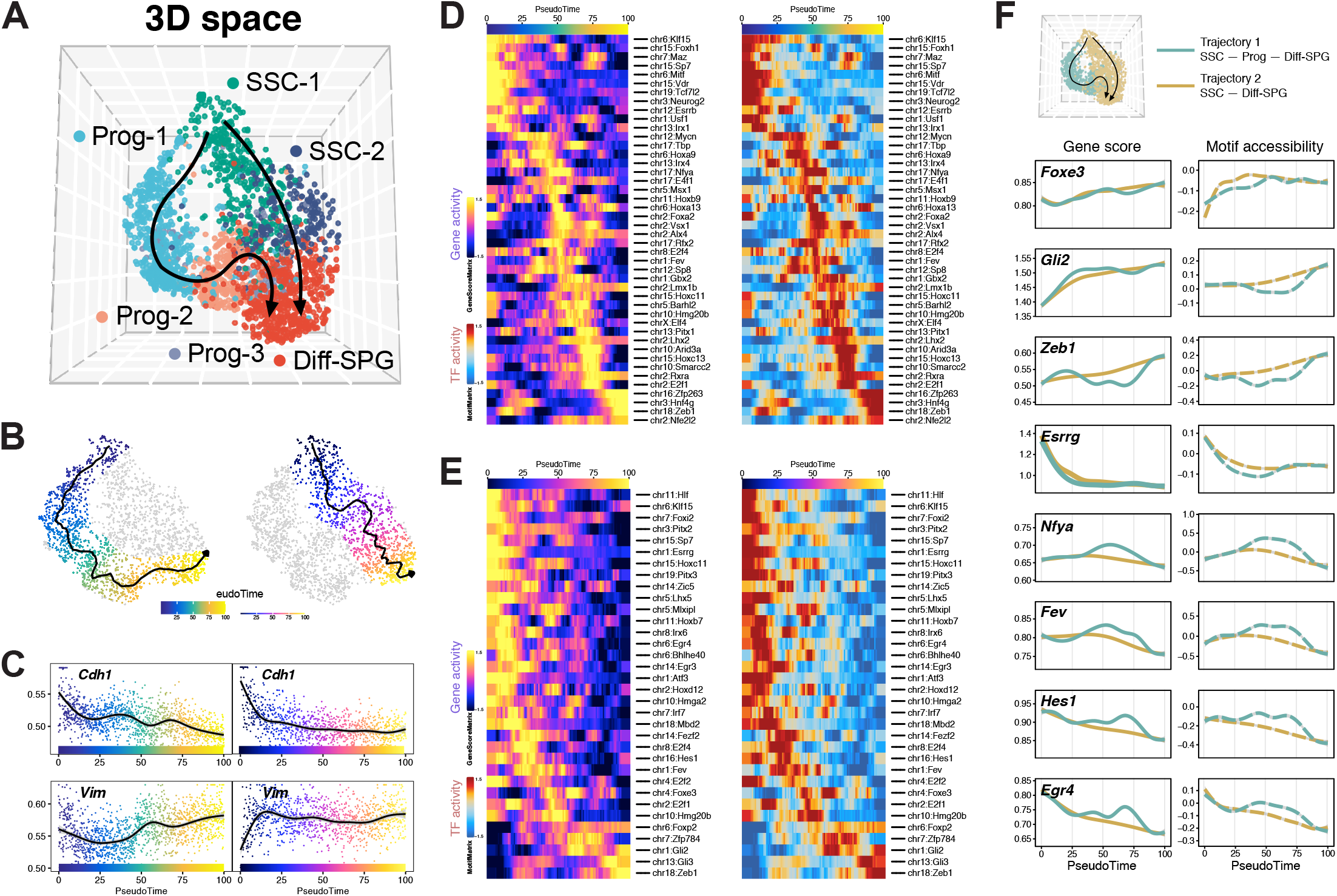
Pseudotime analysis revealed dynamic TF regulation on the EMT-like process in GSC. **A.** De novo identification of two pseudotime trajectories capturing the EMT-like process visualized using UMAP in 3D space. **B.** scATAC-Seq profiles are ordered by pseudotime, corresponding to the epithelial-mesenchymal axis starting from SSC-1 subset to Diff-SPG subset through Prog-1/2/3 subset (left) or SSC-2 subset (right). **C.** Gene score of *Cdh1* and *Vim* in individual cells ordered by pseudotime in Trajectory 1 (left) and 2 (right). **D.** Smoothened heatmap showing dynamic gene score (left) and motif accessibility (right) of indicated TFs along pseudotime for gene-motif pairs in Trajectory 1. **E.** Smoothened heatmap showing dynamic gene score (left) and motif accessibility (right) of indicated TFs along pseudotime for gene-motif pairs in Trajectory 2. **F.** Gene score (left) and motif accessibility (right) of selected TFs identified in D and E ordered by pseudotime. Colored lines correspond to the trajectories shown in 3D UMAP above.

We generated an ordering of single cells (referred to as ‘pseudotime’) along these trajectories (Figure 5B). As expected, gene score of the mesenchymal marker *Vim* varied in a reciprocal gradient to the epithelial marker *Cdh1* across the embedded cells in both trajectories (Figure 5C). We then determined which genes drive the transition in these trajectories through identifying stage-specific factors that co-vary over pseudotime (Supplementary Figure S9). We found prominence of *Hist1* locus genes at the early stage and olfactory receptors at the late stage of pseudotime. Other candidates in the intermediate stage include several genes with known roles in EMT, including *Macrod2, Dock1, Ank2* and *Cdh13.* For instance, overexpression of HIST1H1A could upregulate several EMT regulators[50] and MACROD2 overexpression could reverse EMT[51].

We then tried to identify critical TFs involved in the EMT-like process. chromVAR was inherently based on TF motifs and could not provide a one-to-one correspondence between binding motifs and particular TF proteins, because TFs of the same family often share a similar motif. We reasoned that increasing the abundance of an activator TF will increase its TF activity, or its effect on the state of chromatin, which can be reflected by positive correlation between its expression (gene score) and TF activity (chromVAR z-score). Thus, we correlated gene scores of a TF across pseudotime to its corresponding TF z-score to uncover high confidence regulators. This revealed TFs enriched at the EL-state or ML-state of each trajectory (Figure 5D to F). The EL-state featured ESRRG, EGR4 and SP/KLF family TFs. EGR4 regulates spermatogenesis via several testicular cell types including germ cells [52] and loss of EGR4 leads to premature germ cell death and infertility [53]. ESRRG is a member of a nuclear receptor (NR) superfamily of TFs, which has been proposed as a candidate gene for involvement in human male fertility[54]. Conversely, known EMT regulators are enriched in the ML-state. For instance, GLI2 has been reported to cooperate with ZEB1 for transcriptional repression of CDH1 expression in melanoma cells, while FOXE3 regulates EMT in lens epithelial cells[55].

This analysis also revealed differential TF binding dynamics in each branch, suggesting that a distinct set of TFs cooperatively regulates GSC differentiation and EMT-like transition in each trajectory. Several TFs showed a decreasing trend in Trajectory 1 but showing a transient increase in Trajectory 2, with some of them having known roles in both EMT and male reproduction (Figure 5F). For example, NFYA is overexpressed in some cancers and its motif is related to meiosis-specific gene regulation[56–58]. *Hes1* is expressed in prepubertal spermatogonia in mouse and SSEA+ human SSCs [59, 60]. It can also promote EMT via the AKT pathway in cancers[61]. Lastly, FEV, a solid tumor oncogene, is also upregulated in active-phase testes for cell differentiation[62, 63].

Taken together, our results have provided comprehensive insights into chromatin accessibility dynamics and the distinct TF regulatory programs in GSC undergoing an EMT-like process.

### Epithelial-like and mesenchymal-like-state-specific regulatory topics and enhancers in GSC

We then sought to explore gene enhancer usage in GSC. First, we identified the top 500 differentially accessible regions (DARs) within each cluster. Then, we located the enhancer regions by considering chromatin state revealed by chromHMM using recent ChIP-seq profiling of Id4-GFP spermatogonia[64]. Following this, we uncovered potential stage-specific enhancers of each cluster (Supplementary Figure S10). For instance, the enhancer located in *Zeb1* was the most accessible in Diff-SPG, in accordance with its ML-state. Other potential enhancer regions resided in *Egfr, Foxc1, Dnmt3a, Chd7* and *Cd9* loci, which all have known functions in spermatogonial regulation or spermatogenesis.

The EL-state contained subsets of cells from both SSC-1 and Prog-1, which could not be definitively resolved in the above cluster-oriented analysis. Therefore, we used cisTopic to explore substructure within cell populations associated with a continuous EMT-like process, which is one of the best methods for dynamic populations [44]. Louvain clustering using the regulatory topics identified (Figure 6A) captured similar separation of cell states as shown in SnapATAC results (Figure 4C). Among the regulatory topics, Topics 10, 12, 14 and 16 were most prominent in Prog-1, while Topics 1, 8, 15 and 19 were less enriched in Prog-1 (Figure 6B, Supplementary Figure S11A and B). To gain insight into the function of genes associated with distinct regulatory topics, we performed functional enrichment analysis using GREAT analysis. Topics 10 and 16 were related to stem cell maintenance and division, while Topics 8 and 15 showed direct association with EMT (Supplementary Figure S11C). We then identified topics enriched in EL-state (SSC-1 and Prog-1) and ML-state (Diff-SPG). Three topics (Topics 7, 18 and 22) contributed to the segregation of EL-state, and three topics (Topics 6, 11 and 17) contributed more in the ML-state (Figure 6B and C). GREAT analysis revealed the GO terms enriched in mesenchymal-like topics are associated with NF-kB pathway and response to retinoic acid, which is consistent with the IPA analysis of ML-GSC enriched genes and differentiating spermatogonia, respectively (Figure 6D). Epithelial-like topics were associated with stem cell maintenance and Wnt signaling pathway (Figure 6D). Interestingly, more mesenchymal-like topic regions were located in active enhancer regions (10.3-13.4%), compared with epithelial-like topic regions (4.9-5.0%) (Figure 6E). We then determined specific enhancers in EL- and ML-states (Figure 6F). For example, epithelial-like topic regions included active enhancers located in *Fgfr1* and *Usf1,* which is related to undifferentiated spermatogonia and GSC maintenance[65, 66]. The enhancer region located in *Cdh13* was also more accessible in EL-state, which is consistent with pseudotime analysis. Mesenchymal-like topic regions contained enhancer regions located in *Notch1* and *Mmp9*, which are associated with EMT.

**Figure 6.**
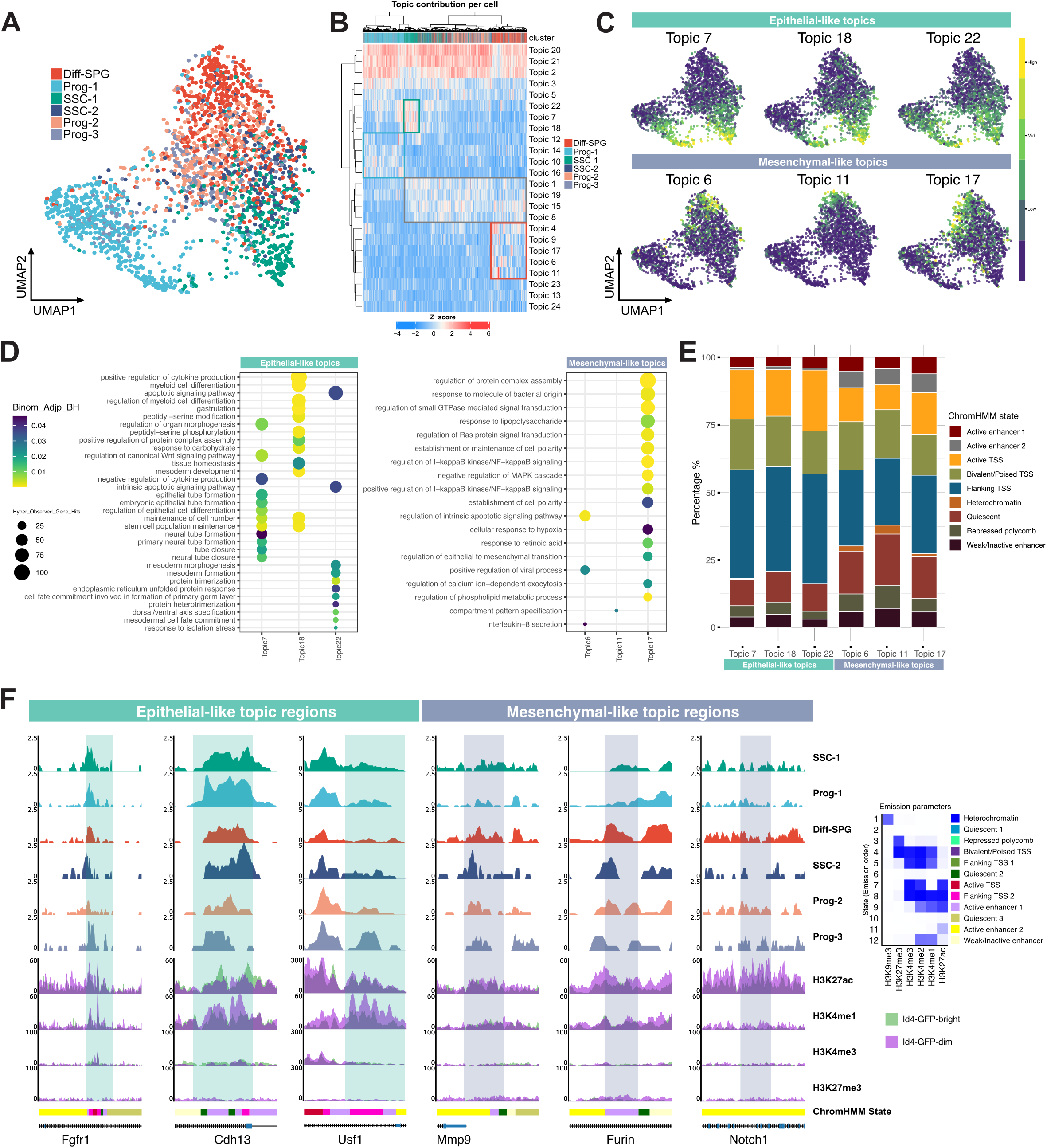
Regulatory topics identified by chromatin accessibility associated with E-M heterogeneity. **A.** cisTopic cell UMAP colored by the SnapATAC cluster assignment. **B.** Heatmap showing Topic-cell enrichment revealed by cisTopic analysis. **C.** cisTopic cell UMAP color-coded by the normalized topic score of selected EMT associated topics. **D.** GREAT analysis of regions included in epithelial-like and mesenchymal-like topics. **E.** Genomic distribution of epithelial-like and mesenchymal-like topic regions revealed by ChromHMM. **F.** Aggregated scATAC-seq profiles showing three regions from epithelial-like topics and three regions from mesenchymal-like topics. ChromHMM tracks indicate chromatin-state based on histone modification profile from Id4-GFP-bright and Id4-GFP-dim spermatogonia (GSE131656). Emission profile (right) from a 12-State model is based on the five histone modifications studied.

Many enhancer elements are not located directly adjacent to the gene whose expression it regulates but reside far from the transcription start sites. Besides, relationships between enhancers and target genes are not always one-to-one. Therefore, it is still challenging to connect enhancer elements to their target genes[67]. Leveraging a recently described strategy, Cicero[68], we linked distal regulatory elements to their prospective genes via patterns of co-accessibility (Figure 7A). First, we were able to interconnect 26,927 scATAC-Seq peak pairs at a co-accessibility score cut-off of 0.25, in which 5,082 peak pairs linked a distal regulatory element to a promoter. Of these, 1,037 links included distal elements that fell within active enhancer regions, which might represent putative enhancer-promoter interactions. GO analysis of the target genes revealed significant enrichment of EMT regulation (Figure 7B). For example, regions marked by both H3K4me1 and H3K27ac were connected to the promoters of *Notch1* and *Bmp7,* suggesting putative enhancers regulating their expression during the EMT-like process (Figure 7C). Other examples included enhancers for *Spry4* and *Lin28a,* which are both highly expressed in undifferentiated spermatogonia, including SSC (Figure 7C)[69, 70].

**Figure 7.**
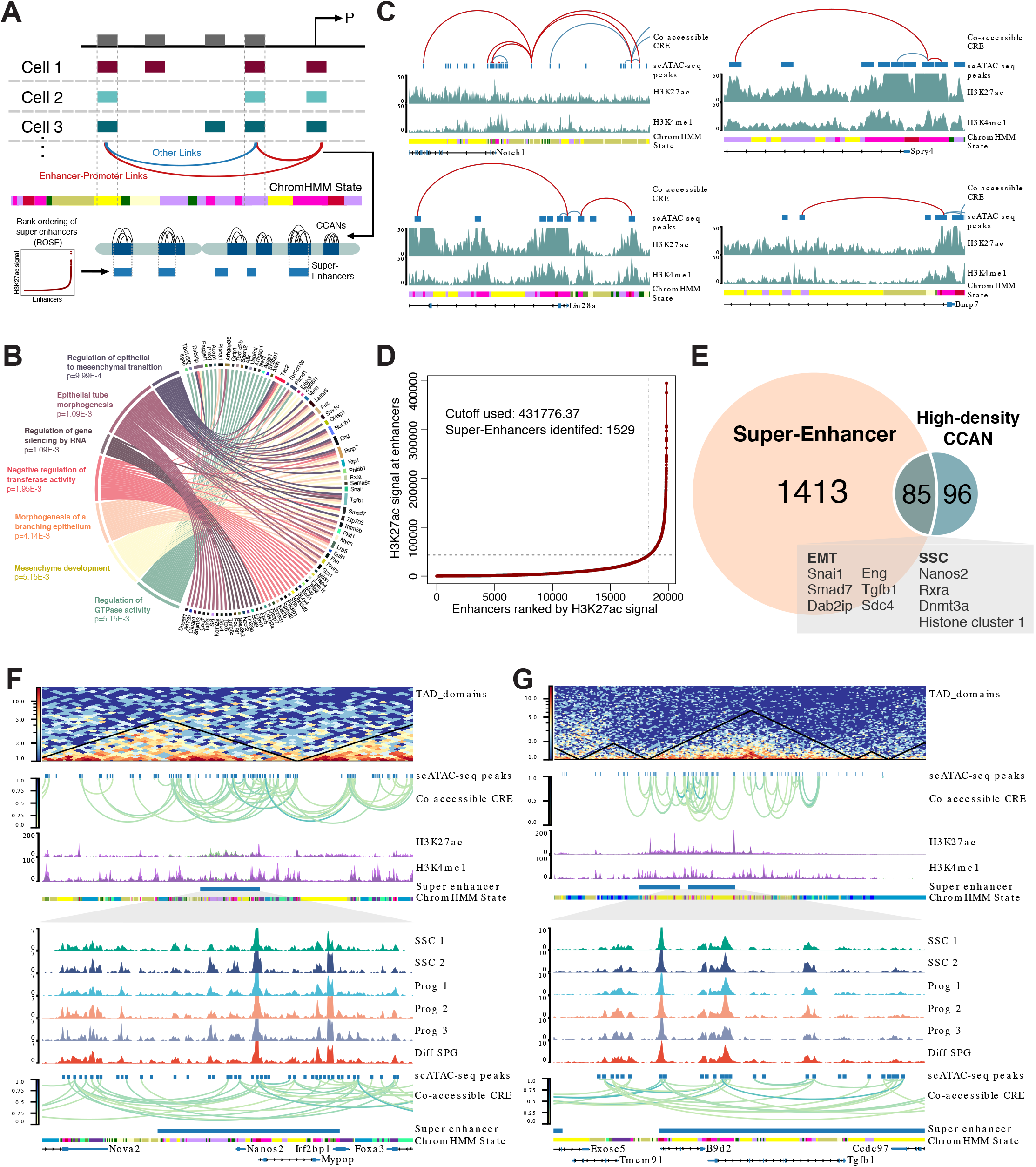
Cis-coaccessibility analysis revealed putative enhancers in GSC. **A.** The schematic diagram illustrating the approach to link enhancer to target gene. **B.** The Circos plot represents significantly enriched pathways and GO terms for biological process associated with putative enhancer-promoter links identified. Outside the circle, genes and significantly enriched pathways together with GO terms are indicated. **C.** Inferred links for *Notch1, Spry4, Lin28a, Bmp7.* Red and blue lines indicate enhancer-promoter links and other links respectively. ChromHMM tracks indicate chromatin-states related to Figure 4. **D.** Call of super-enhancers (SEs) by Rank ordering of super-enhancers (ROSE), which takes into account enhancer ranks by H3K27ac signals from Id4-GFP-bright spermatogonia ChIP-seq profile (GSE131656). **E.** Venn diagram showing significant overlapping of SEs identified from ROSE and high density cis co-accessibility networks (CCANs). **F.** Aggregated scATAC-seq profiles showing SE regions near *Nanos2* with inferred TAD domains[72] and co-accessible CREs. **G.** Aggregated scATAC-seq profiles showing SE regions near *Tgfb1* with inferred TAD domains[72] and co-accessible CREs.

Further, we identified cis-co-accessibility networks (CCANs) using Cicero, which can inform us about modules of coregulated chromatin in the genome. We focused on the 181 regions with an exceptionally large number of significant co-accessible peak links (high-density CCANs, greater than 10 links), which incorporated 3626 sites. GREAT analysis associated the peaks contained in these high-density CCANs to 1181 genes and revealed significant enrichment in EMT regulation (Binom FDR Q-Value, 2.80e-79). We further examined the relationship of high-density CCANs and super-enhancers, which are clusters of enhancers highly enriched in histone H3K27 acetylation [71]. We found high-density CCANs significantly overlapped the 1529 super-enhancers identified by ROSE based on SSC H3K27ac mark (p <2.489e-61, hypergeometric test) (Figure 7D and E). Accordingly, genes within high-density CCANs were strongly enriched in super-enhancer associated genes (p < 8.339e-09, hypergeometric test, representation factor = 1.5). Focusing on individual loci, we observed key EMT and spermatogonia related genes that exhibited patterns of coordinated regulation, which tended to occupy the same topologically associated domain (TAD) in spermatogonia[72]. We found super-enhancer regions located at *Nanos2* and *Tgfb1* loci. In accordance with the role of *Nanos2* in SSC maintenance, several peaks in the super-enhancer region were more accessible in SSC-2 subset (Figure 7F). Consistent with the role of TGF-β signaling in the EMT-like process, we found that the peaks at *Tgfb1* locus were less active in GSCs at EL-state (SSC-1 and Prog-1) (Figure 7G). We also identified super-enhancers at *Dab2ip* and histone cluster 1 loci. *Dab2ip* facilitates spermatogonial differentiation and overexpression of certain histones promotes EMT [73] [50] (Supplementary Figure S12).

Taken together, scATAC-Seq revealed that the EMT-like process is dynamically orchestrated by distal enhancers and enhancer-to-gene interactions.

### Potential target of G9a regulation in GSC

Having established the chromatin states associated with EMT-like transition in GSC using scATAC-seq, we sought to assess the role of G9a in epigenetic control through analyzing the datasets of G9a inhibition. First, we generated synthetic pseudo-bulk datasets by merging data from cells in control and UNC0638-treated groups and identified DARs between these two conditions. We found chromatin accessibility of genes inducing EMT *(Tgfbr1* and *Klf9*) were downregulated after G9a inhibition while genes suppressing EMT (*Cdh13* and *Fgf21*) showed DARs with increased accessibility (Figure 8A). Examining the distributions of DARs in chromHMM chromatin states revealed that regions gained accessibility showed a higher percentage of peaks mapped to quiescent chromatin states and heterochromatin (Figure 8B). Theser regions may be initially bound with inhibitory histone modification such as H3k9me2/3, which is in line with the role of G9a’s role in heterochromatin formation. Conversely, regions with reduced accessibility were more enriched for super-enhancers, suggesting repression of gene transcription (Figure 8C).

**Figure 8.**
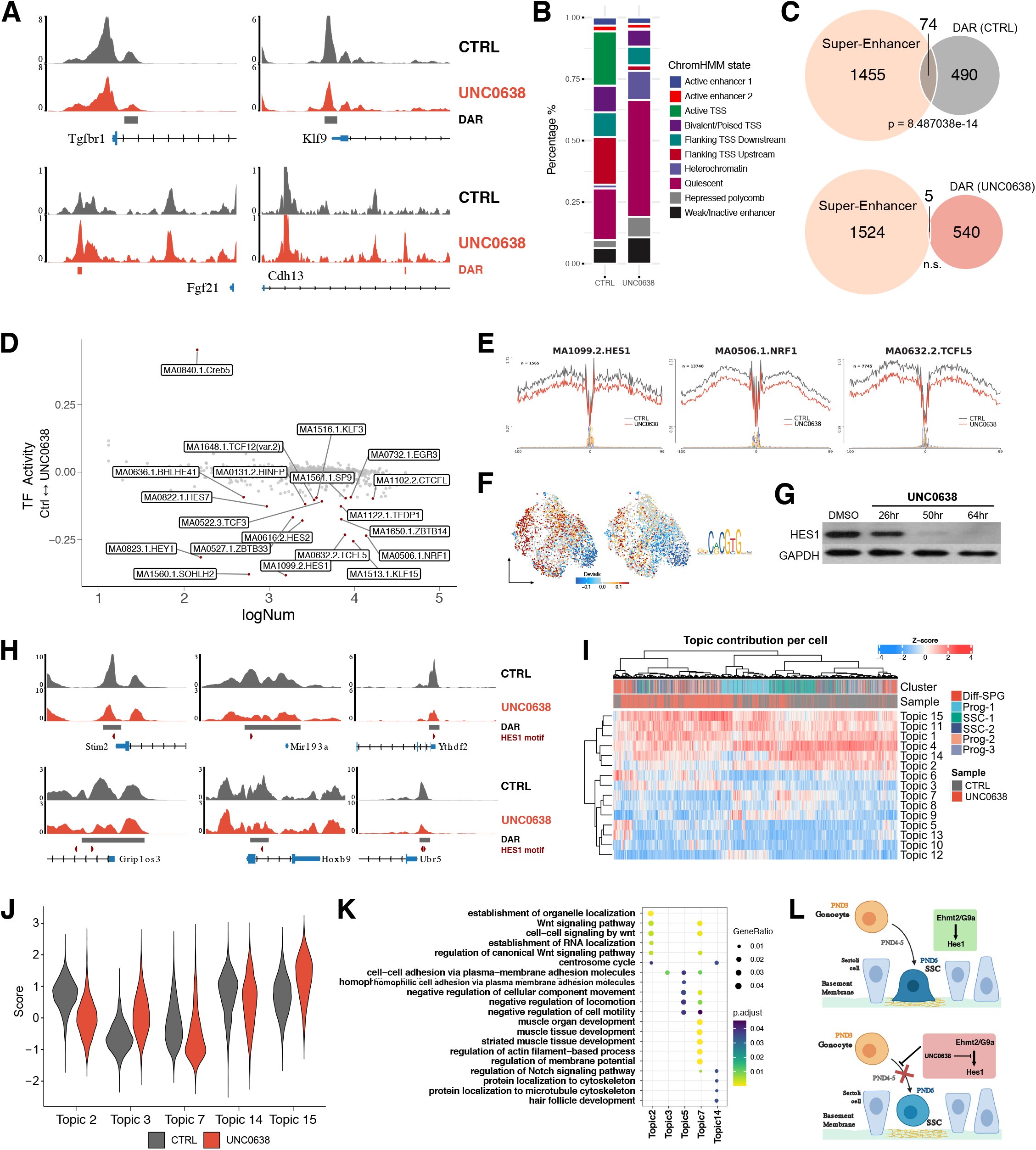
Gene expression and chromatin accessibility changes associated with G9a inhibition. **A.** Normalized pseudo-bulk accessibility tracks showing DARs of selected EMT associated genes between control and UNC0638 treatment group. **B.** Genomic distribution of differentially accessible regions (DARs) of control (upper panel) and UNC0638 treatment (lower panel) group revealed by ChromHMM. **C.** Venn diagram showing overlapping of super-enhancers from Figure 4D and DARs of control (top) and UNC0638 treatment (bottom) group. **D.** Scatter plot showing the transcription factor (TF) activity dynamics between control and UNC0638 treatment group. Y-axis represents the differences in TF activity and the x-axis indicates the number of cleavage events (log10 transformed). TFs with significant differential activity identified by HINT are colored as red (adjusted p-value < 0.05 and the number of cleavage events > 150). **E.** Transcription factor footprints (average ATAC-seq signal around predicted binding sites) for selected TFs. Logo of underlying sequences is shown below. **F.** Motif activity of HES1 in control and UNC0638-treated GSCs shown in UMAP. Logo of HES1 motif is shown on right. **G.** Western blotting of HES1 in control (DMSO) and UNC0638-treated (2 μM) cells at 26, 50 and 64 hours after treatment. **H.** Normalized pseudo-bulk accessibility tracks showing DARs with HES1 binding motif between control and UNC0638 treatment group. **I.** Heatmap showing Topic-cell enrichment combining both the control and UNC0638 treatment groups revealed by cisTopic analysis. **J.** Violin plot of the normalized topic score of topics that lose accessibility (Topic 2, 7 and 14) or gain accessibility (Topic 3 and 15) after UNC0618 treatment. **K.** GO analysis of regions including in topics with differential accessibility between control and UNC0638 treatment group. **L.** Schematic diagram illustrating that the migration of gonocyte towards the basement membrane to establish SSC pool requires G9a to regulate *Hes1* (top). This process is inhibited upon the addition of G9a inhibitor UNC0638 (bottom).

Accessibility at regulatory sites is driven by TF binding. Therefore, a comparison of the TF binding activity based on differential footprinting will allow us to reveal molecular mechanisms of G9a-mediated regulation of gene expression. Surprisingly, we observed most TFs exhibited significant loss of activity after UNC0638 treatment with scHINT analysis with the exception of CREB5 (Figure 8D and E) [74]. The most prominent factors are members of the Hes/Hey family. Among them, *Hes1* was also identified to be associated with the EMT-like trajectory in pseudotime analysis. At single cell level, in accordance with the EMT suppression, the proportion of mesenchymal-like cells (Diff-SPG) decreased after UNC0638 treatment (Figure 4D). Conversely, we observed an increased proportion of cells mainly in Prog-2. This raised the possibility that G9a might have a more profound effect on Trajectory 1, which is consistent with a higher activity of HES1 motif in Prog-2 along Trajectory 1 (Figure 8F). Concordantly, HES1 motif activity in Prog-2 substantially decreased after G9a inhibition. We further examined HES1 protein level and found that it drastically decreased upon G9a inhibition over time and was nearly depleted after 64 hours (Figure 8G). *Hes1* mRNA level also showed a mild decrease trend after UNC0638 treatment from bulk RNA-Seq. These results suggested that G9a might regulate GSC EMT-like process through HES1. Next, we tried to identify genes that may be the direct regulatory targets of HES1 underlying G9a regulation. We reasoned that target genes would be associated with HES1 motif-containing DARs with reduced accessibility after G9a inhibition. Interestingly, we found several putative targets that were directly associated with EMT regulation, including *Ythdf2*, *Stim2, Grip1, Hoxb9, Ubr5,* and *Mir193a* gene locus (Figure 8H) [75–80].

Since scHINT analysis only identified CREB5 with higher TF activity after G9a inhibition, we next sought to test whether a different approach, i-cisTarget, would reveal more potential targets with increased activity. We analyzed the DARs with higher accessibility after UNC0638 treatment. In line with the methyltransferase activity of G9a, we found a significant enrichment (NES = 8.54143) for H3K9me3 regions, which gained accessibility after G9a inhibition. Moreover, there is an enrichment (NES = 6.62392) for REST, which is known to negatively regulate EMT through transcriptional repression [81].

We next used cisTopic to compare the control and UNC0638 treatment group (Figure 8I). cisTopic identifies co-accessible regulatory regions (cis-regulatory topics) for each cell by clustering all scATAC-Seq peaks. This identified three “loss of accessibility topics”, Topic 2, 7 and 14 (Figure 8J). Topic 2 is highly enriched for genes associated with Wnt pathway; Topic 7 is related to actin-filament regulation while Topic 14 is related to Notch signaling pathway (Figure 8K). These pathways have been suggested to be necessary for EMT[82–84]. On the other hand, we identified two topics that represent an increase in accessibility (Figure 8J). Topic 3 contains regions significantly associated with cell-cell adhesion while Topic 15 is associated with general transcriptional regulation (RNA splicing) (Figure 8K).

Collectively, our transcriptomic and epigenetic analysis allow the identification of the underlying mechanism of G9a in the regulation of EMT-like process and its potential role in germ cell migration through HES1 (Figure 8L).

## Discussion

Analysis of several gene expression profiling studies have indicated that E/M heterogeneity exists in spermatogonia in vivo. However, the biological relevance of this phenomenon is yet to be determined. In this study, we found the regulation of G9a in spermatogonia may involve an EMT-like process. Following this, we demonstrated the epithelial-mesenchymal plasticity of GSCs, as we identified two interconvertible subtypes of GSCs with completely different cellular and molecular properties when cultured on Matrigel. We further showed that GSC migration is regulated by EMT-related mechanisms.

Despite the known role of G9a as an essential epigenetic regulator during spermatogenesis, such studies have been predominantly limited in scope to male germ cell meiotic progression [85]. Existing data indicated that G9a exerts a transcriptional repression function dependent on its HMTase activity that mediates H3K9 methylation[86]. G9a might cause repression of regulators which in turn controls the EMT-like process in GSC. A plausible candidate is *Cdh1*. We showed that G9a inhibition by UNC0638 led to an increase in CDH1, possibly due to the removal of H3K9me2 on its gene regulatory elements. In line with this, G9a could cause transcriptional repression of *CDH1* in cancers [87, 88]. Loss of CDH1 results in the liberation of β-catenin that in turn induces expression of EMT-TFs [89], which might contribute to the downregulation of EMT genes after G9a inhibition. Conversely, G9a inhibition downregulated HES1 in both protein level and TF activity in GSC, which has been reported to promote EMT and cell migration [90]. The underlying regulation can be explained by the function of G9a as a coactivator, since it has been reported that G9a activates the Notch signaling pathway by regulating *Notch1* and *Hes1* in cancer progression of melanoma[91]. Another possibility is that G9a might promote AKT activity which in turn upregulates HES1[61, 92]. Interestingly, we observed increased chromatin accessibility of *Notch1* and *Akt* in mesenchymal-like regulatory topic regions. implying their potential role in the EMT-like process. Yet, further investigation is needed to uncover the mechanisms of G9a-dependent regulation of the EMT-like process in GSC and germ cell migration.

Although genes related to germ cell migration such as the TF*RhoxlO* and actin disassembly factor *Aip1* have been reported[73, 93], mechanisms that regulate germ cell migration are largely unknown due to the lack of an effective cellular model. In this study, we established a manageable Matrigel-based culture system to recapitulate the EMT-like process and study cell migration in GSC in vitro. Matrigel is a reconstituted cell culture matrix able to control stem cell fate by influencing growth, differentiation and migration[94]. Our results revealed that high concentrations of Matrigel coating induced an EMT-like process as it promoted the formation of flat-shaped GSC colonies and mesenchymal-like cells. This phenotypic change was mainly mediated by TGF-β signaling as evident by RNA-seq analysis. In the SSC niche, ECM proteins contributing to the basement membrane include type IV collagens, laminin, heparan sulfate proteoglycan (HSPG) and entactin, many of which are contained in Matrigel[95]. Among them, HSPGs have been demonstrated to regulate several major signaling pathways, including TGF-β, fibroblast growth factor (FGF), Wnt and Hedgehog (Hh) pathways. It has been shown that FGFs can be immobilized and concentrated on the HSPG-rich basement membrane in testis[5] and FGFs can also augment TGF-β1-induced EMT in mammary epithelial cells[96]. Therefore, it is possible that increased deposition of HSPGs ensures an enriched pool of bioactive growth factors (e.g. TGF-β1 and FGFs) in the accumulating matrix to promote the EMT-like process in GSC.

Furthermore, our culture system provides a valuable platform to study the dynamics of germ cell migration in vitro. We showed that GSC migration is enhanced in the mesenchymal-like population. Further, we found that the gene expression signatures of ML-GSC are concordant with a recently discovered migratory pro-spermatogonia (ProSG) subset, I-ProSG[37], both of which showed upregulation of migratory genes. This suggested a shared mechanism of GSC migration in vivo and in vitro.

To our knowledge, our study is the first to dissect the complex epigenetic regulatory networks of GSCs at single cell resolution. Relatively little is known about chromatin accessibility dynamics during SSC differentiation. Previous ATAC-Seq analysis of spermatogenesis revealed that open chromatin in differentiated spermatogonia (KIT+ cells) is similar to that in undifferentiated spermatogonia (THY1+ cells) and differentiation only involves chromatin closure without de novo formation of accessible chromatin[97]. However, bulk ATAC-Seq may mask true biological signals due to the proportions of the different cell states. We also provide evidence at the single cell level that GSC differentiation can proceed by different routes before converging on the differentiated state. Instead of a linear path, the differentiation pathway of our data resembles a “doughnut” shape, which mirrors the findings from scRNA-seq on human germ cells[98].

Regarding the relationship between GSC differentiation and the EMT-like process, our scATAC-Seq suggested that SSC-1 and Diff-SPG clusters coincide with EL- and ML-like states, respectively. We found decreased DMRT1 TF activity in the mesenchymal-like subset Diff-SPG. DMRT1 is one of the top motifs enriched in human SSC-specific ATAC-seq peaks and loss of DMRT1 causes spermatogonia to enter meiosis[49, 59]. This suggested that SSCs initiate differentiation when entering the mesenchymal-like state. Supporting this, decreased DMRT1 TF activity was accompanied by increased gene scores of differentiation genes. This observation resembles the findings in embryo development, in which EMT has been depicted as a process of differentiation, with gradual loss of potency[99]. Therefore, we speculate that the differentiation program is coupled with an EMT-like process in SSCs. Our results provide a clue to the link between EMT and SSC differentiation. However, the functional implications of our observation requires further validation using transplantation assay in the future.

scATAC-Seq analysis also brings the possibility to learn epigenetic regulation related to E/M plasticity. First, we observed gene scores and TF activity of several canonical EMT regulators displaying distinct patterns across the cell states, the most significant of which was the TF ZEB1. Multiple analyses suggested it increased in activity during the mesenchymal-like state, suggesting a role in the GSC EMT-like transition. This is in line with a recent study showing *Zeb1* as a potential MET suppressor in mouse SSC[100]. We also identified gene signatures (e.g. *Smarcc1, Bach1* and *c-Fos)* that were not typically considered as canonical EMT regulators but have been previously implicated in EMT. Second, based on predicted activity of these canonical EMT genes, cells can be regarded as progressively transitioned throughout the E/M spectrum. When assessing dynamics of chromatin accessibility throughout the pseudotemporal trajectories, we further identified novel regulators of the GSC EMT-like process, such as *Hes1*. Third, scATAC-Seq also reveals cis-regulatory elements associated with E/M plasticity. Using cisTopic, we were able to identify epithelial- or mesenchymal-associated regulatory topics. Annotating these putative regulatory elements suggested signals from multiple cues orchestrate the proper execution of EMT/MET cycles in GSC. For example, the Wnt signaling pathway showed activation, whereas NF-kB and TGFβ activity decreased in epithelial-like clusters. These regulatory elements represent interesting candidates for future studies.

Using our culture model, we have shown that ATAC-Seq performed on heterogeneous samples at the single-cell level can reveal important dynamic regulatory regions across the EMT-like process. Moreover, combined with the analysis of histone modifications, our data allowed us to identify putative enhancer regions more accessible in different cell states. scATAC-Seq also provides a unique opportunity to identify tightly coordinated co-accessible regulatory elements. We highlighted several potential enhancer-promoter interactions and super-enhancer regions associated with key EMT regulators and SSC-related genes. Our dataset, which represents an initial characterization of genome-wide chromatin accessibility and candidate distal regulatory sites in GSC EMT-like process, should be a valuable resource for investigators interested in male germ cell biology, as well as EMT in general.

Of note, our study focused on chromatin accessibility and did not include scRNA-Seq analysis of GSCs. Recently, multimodal single cell profiling (“multi-omics”) has greatly improved our ability to detect unique cell types and states. Joint data of scATAC-Seq and scRNA-Seq from the same cell showed that they are comparable for determining cell identity, suggesting cells coordinate chromatin structure with transcription. Nevertheless, some cell states may not reflect equally in both profiles. Future studies would benefit from joint profiling to provide additional information that further elucidates E/M heterogeneity.

## Conclusions

Our scATAC-Seq analysis redefines cellular heterogeneity of the undifferentiated population driven by specific EMT-TFs. Our study thus takes a significant step forward in revealing the complex picture of male germ cell development. The concept that neonatal spermatogonial development may also involve an EMT-like process can provide a valuable model to shape future research.

## Methods

### SSC isolation and derivation of GSC long-term culture

Oct4-EGFP transgenic mice (B6; CBA-Tg(Pou5f1-EGFP)2Mnn/J, Stock no.: 004654) were acquired from The Jackson Laboratory and maintained in CUHK Laboratory Animal Services Center[101]. Spermatogonial stem cells from testes of Oct4-GFP transgenic mice at PND5.5 were purified using the method described previously[102]. Cells were resuspended in medium with a density of 100,000 cells per well of a 24 well tissue culture plate, either on MEF layer or Matrigel-coated well.

### scATAC-Seq library preparation and data analysis

Cell tagmentation was performed according to SureCell ATAC-Seq Library Prep Kit (17004620, Bio-Rad) User Guide (10000106678, Bio-Rad). In brief, washed and pelleted cells were lysed with the Omni-ATAC lysis buffer. Nuclei were counted and examined under microscope to ensure successful isolation. Same number of nuclei were subjected to tagmentation with equal ratio of cells/Tn5 transposase to minimize potential batch effect. Nuclei were resuspended in tagmentation mix and agitated on a ThermoMixer for 30 min at 37 °C. Tagmented nuclei were loaded onto a ddSEQ Single-Cell Isolator (Bio-Rad). scATAC-Seq libraries were prepared using the SureCell ATAC-Seq Library Prep Kit (17004620, Bio-Rad) and SureCell ATAC-Seq Index Kit (12009360, Bio-Rad). Libraries were sequenced on HiSeq 2000 with 150 bp paired-end reads. Sequencing data were processed using the Bio-Rad ATAC-Seq Analysis Toolkit. Downstream analysis was performed with SnapATAC[43], cisTopic [103] and ArchR[104].

### Statistical analysis

Statistical analysis. Assessment of statistical significance was performed using two-tailed unpaired t-tests, one-way ANOVA with Tukey multiple comparisons tests or Chi-squared tests. Statistical analysis was performed using GraphPad Prism v8. Associated P values are indicated as follows: *P < 0.05; **P< 0.01; ***P < 0.001; ****P < 0.0001; not significant (ns) P > 0.05.

### Ethics approval and consent to participate

All the animal experiments were performed according to the protocols approved by the Animal Experiment Ethics Committee (AEEC) of The Chinese University of Hong Kong (CUHK) and followed the Animals (Control of Experiments) Ordinance (Cap. 340) licensed from the Department of Health, the Government of Hong Kong Special Administrative Region.

## Consent for publication

Not applicable.

## Availability of data and materials

The datasets generated and/or analysed during the current study are available in the Gene Expression Omnibus (GEO) repository, under accession numbers GSE159412 and GSE159414.

## Authors’ contributions

JL, HCS, RMH and TLL conceived and designed the study. JL and HCS performed experiments. JL, HCS, RMH and TLL analyzed and interpreted data. JL, HCS, RMH and TLL wrote the paper. JL, HCS, ACSL, WTL, JKWN, THTC, MYC, DYLC, HQ, WYC, RMH and TLL reviewed and edited the manuscript. All authors read and approved the final manuscript.

## Acknowledgements

We thank Tommy Lo in Bio-rad for scATAC-Seq experimental assistance. We also thank the SBS Core Laboratories in The Chinese University of Hong Kong for technical support.

