## Supplemental methods and figure legends for "scATAC-Seq reveals epigenetic heterogeneity associated with an EMT-like process in male germline stem cells and its regulation by G9a"

### Supplementary Methods

#### Animals

All the animal experiments were performed according to the protocols approved by the Animal Experiment Ethics Committee (AEEC) of The Chinese University of Hong Kong (CUHK) and followed the Animals (Control of Experiments) Ordinance (Cap. 340) licensed from the Department of Health, the Government of Hong Kong Special Administrative Region. All the mice were housed under a cycle of 12-hour light/dark and kept in ad libitum feeding and controlled the temperature of 22-24°C. Oct4-EGFP transgenic mice (B6; CBA-Tg(Pou5f1-EGFP)2Mnn/J, Stock no.: 004654) were acquired from The Jackson Laboratory and maintained in CUHK Laboratory Animal Services Centre[1].

#### SSC isolation and derivation of long-term culture

Spermatogonial stem cells from testes of Oct4-GFP transgenic mice at PND5.5 were purified using the method described previously[2]. Isolated seminiferous tubules were digested with 1 mg/ml type 4 collagenase (Gibco), 1 mg/ml hyaluronidase (Sigma-Aldrich) and 5 µg/ml DNase I (Sigma-Aldrich) at 37°C for 20 min with occasional shaking. The suspension was passed through a 40- µm strainer cap (BD Falcon) to yield a uniform single cell suspension. After incubation in staining buffer (PBS supplemented with 1% FBS, HEPES, glucose, pyruvate and penicillin-streptomycin) with APC anti mouse CD117 ckit antibody (553356, BD Biosciences) at 4°C for 30 min, Oct4-GFP+/KIT- cells were collected with a BD FACSAria Fusion Flow Cytometer (BD Biosciences). Primary SSCs were then cultured on mouse embryonic fibroblast layer (150,000 cells per well of a 24 well tissue culture plate). GSC medium (Supplementary Table S6) was used for GSC maintenance. GSCs were passaged by treating with 0.25% trypsin (Invitrogen) for two minutes and adding fresh medium to stop trypsinization. After centrifugation at 1000 rpm for 5 minutes, the cell pellet was resuspended in medium with a density of 100,000 cells per well of a 24 well tissue culture plate, either on MEF layer or Matrigel-coated well.

#### Western blot

GSC cultured on MEF feeder were treated with 2 mM UNC0638 for 26, 50 and 64 hours. Cells were harvested and FACS sorted to remove MEF feeder cells. Cell lysate was separated by SDS/PAGE, transferred to Immobilon-P membrane (Millipore), and incubated with primary antibodies and HRP-conjugated secondary antibodies (Supplementary Table S7). Immunoreactive bands were detected using Western Lightning Plus-ECL (Perkin Elmer). The antibodies used are listed in Supplementary Table S2.

#### In vivo G9a inhibitor UNC0638 treatment

Oct4-EGFP transgenic mice were used. Treatment with G9a inhibitor was performed by oral administration of UNC0638 (Abcam) diluted in 1% tragacanth (G1128, Sigma) with a dosage of 3 mg/kg daily on PND3-5. DMSO diluted in 1% tragacanth was used as a control. For each litter, at least one male was used for drug treatment and at least one male was used as control. On PND6, the mice were euthanized and their testes were collected. After being fixed in 4% (v/v) paraformaldehyde for 2 hours, 10 μm cryosections of neonatal mouse testes were prepared as described[3].

### Bulk RNA-Seq

Low input RNA-Seq library preparation was conducted according to the modified protocol of Smart-seq2[4]. In brief, the mixture of 1 μl of 10 mM oligo-dT30VN primer (5'-AAGCAGTGGTATCAACGCAGAGTACT30VN-3'), 1 μl of 10 mM dNTP mix and 10 ng extracted RNA in 5 μl reaction volume was incubated at 72°C for 3 min. Then, 5 μl of the reverse transcription mixture (200U SuperScript II reverse transcriptase, 20U RNase inhibitor, 1x Superscript II first-strand buffer, 5 mM DTT, 1 M Betaine, 6 mM MgCl2, 1 μM TSO (5'- AAGCAGTGGTATCAACGCAGAGTACATrGrG+G-3') was added into the tube for RNA reverse transcription. The tubes containing a total 10 μl mixture were placed in ProFlex PCR System (Thermo Fisher Scientific) and the reaction condition was as follows: 42°C for 90 min, followed by ten cycles of 50°C for 2 min and 42°C for 2 min. The final enzyme inactivation was performed at 70°C for 15 min. After finishing the first-strand reaction, the 2x PCR preamplification mix (1x KAPA HiFi HotStart ReadyMix, 0.1 μM IS PCR primers (5'- AAGCAGTGGTATCAACGCAGAGT-3')) was added. PCR preamplification reaction was performed in ProFlex PCR System by using the following program: 98°C for 3 min, 12 cycles of 98°C for 20 s, 67°C for 15 s and 72°C for 6 min, with the final extension cycle of 72°C for 5 min. Amplified cDNA was purified by using Ampure XP beads (1:1 ratio) according to the manufacturer's manual and was eluted in 15 μl of ddH2O. The yield of amplified cDNA was measured by Qubit3.0 fluorometer. Amplified cDNA with good quality were used for library construction with TruePrep DNA Library Prep Kit V2 for Illumina (Vazyme) following the instruction. The quality of all constructed libraries was assessed by Agilent Bioanalyzer 2100 with High Sensitivity DNA Kit (Agilent Technologies). Qualified libraries were sequenced on Illumina HiSeq2000 to generate 150bp paired-end reads. Raw reads were pre-processed with trim-galore for trimming Illumina adaptor sequences (http://www.bioinformatics.babraham.ac.uk/projects/trim_galore/). Counts were assigned to genes defined by the Ensembl (release 67) annotation using RSEM[5]. Differentially expressed genes were identified with edgeR package[6]. Soft clustering using differentially expressed genes was obtained using the Mfuzz package implemented in R[7]. GO enrichment and pathway analysis was performed using ToppGene Suite[8], Ingenuity Pathway Analysis (IPA) and GSEA[9]. A hypergeometric test with the Benjamini and Hochberg false discovery rate (FDR) was performed using the default parameters to adjust p-value.

### Bulk ATAC-Seq

ATAC-Seq was performed with ML- and EL-like GSCs according to Omni-ATAC protocol using TruePrep DNA Library Prep Kit V2 for Illumina (TD501) with modifications[10]. Briefly, around 100,000 cells were harvested and resuspended in 25 μL ATAC-Resuspension Buffer (RSB) containing 0.1% NP40, 0.1% Tween-20, and 0.01% Digitonin and incubated on ice for 3 min. The cell lysis was washed out with 0.5mL cold ATAC-RSB containing 0.1% Tween-20. After centrifugation at 500g for 10 mins, pellet nuclei were resuspended in 10ul PBS. Nuclei were counted and 50,000 nuclei were mixed in 50 μL of transposition mixture (10 μL 2x TTBL buffer, 5 μL transposase (TTE mix), 16.5 μL PBS, 0.5 μL 1% digitonin, 0.5 μL 10% Tween-20) by pipetting up and down. Tagmentation were performed at 37°C for 30 min in a thermomixer with 1000 RPM mixing. The transposed DNA was purified with Zymo DNA clean and concentrator-5 kit. Subsequently, purified transposed DNA was then amplified by PCR for 7 cycles with TruePrep DNA Library Prep Kit. The quality of all constructed libraries was assessed by Agilent Bioanalyzer 2100 with High Sensitivity DNA Kit (Agilent Technologies). Qualified libraries were sequenced on Illumina HiSeq2000 to generate 150bp paired-end reads. Data were analyzed withATAC-Seq pipeline from nf-core with default setting[11]. Peaks identified by MACS2[12] were used for differential analysis with MAnorm[13].

### scRNA-Seq analysis

Previously published scRNA-Seq datasets were downloaded from GEO (GSE112880 and GSE109049) [14,15]. UMI count tables of each cellular barcode were analyzed using Seurat (Butler et al. 2018). Seurat objects were built by loading UMI count tables using the “Read10X” function. Each sample was filtered and cells with greater than 1000 genes expressed and less than 20% of reads mapped to the mitochondrial genome were retained. After normalization, the top 2000 highly variable genes were selected using the “FindVariableFeatures” function of the Seurat package. Dimensionality of data was reduced by principal component analysis (PCA) (30 components) and visualized with UMAP. Clustering was performed using Louvain algorithm on 30 principal components (resolution = 0.5). After initial clustering, cluster-specific markers were identified with the “FindAllMarkers” function using default parameters and clusters were assigned based on known markers. Only Kit- clusters were included in subsequent analysis. The generated Seurat object was loaded to BioTuring for marker visualization in UMAP plots.

### scATAC-Seq analysis

#### Sample collection

For feeder-free culture group, cells were cultured on Matrigel-coated plates (10 µg/cm2) for 1, 7 and 14 days respectively after being transferred from MEF-dependent culture. Samples at different time points are pooled together in a ratio of 1:2:2 (Day1:Day7:Day14). For G9a inhibition group, cells cultured on MEF feeder were treated with 2 mM UNC0638 for 26, 50 and 64 hours respectively. Samples at different time points are pooled together in equal numbers. For the control group, cells were cultured on the MEF feeder. Cells from each time point were purified using FACS to remove MEF feeder, cell debris and cell aggregate, and pooled (total 120,000 cells for each sample) as mentioned.

#### Cell lysis and tagmentation

Cell tagmentation was performed according to SureCell ATAC-Seq Library Prep Kit (17004620, Bio-Rad) User Guide (10000106678, Bio-Rad) and the protocol based on Omni-ATAC was followed (Corces et al. 2017). In brief, washed and pelleted cells were lysed with the Omni-ATAC lysis buffer containing 0.1% NP-40, 0.1% Tween-20, 0.01% digitonin, 10 mM NaCl, 3 mM MgCl2 and 10 mM Tris-HCl pH 7.4 for 3 min on ice. The lysis buffer was diluted with ATAC-Tween buffer that contains 0.1% Tween-20 as a detergent. Nuclei were counted and examined under microscope to ensure successful isolation. Same number of nuclei were subjected to tagmentation with equal ratio of cells/Tn5 transposase to minimize potential batch effect. Nuclei were resuspended in tagmentation mix, buffered with 1×PBS supplemented with 0.1% BSA and agitated on a ThermoMixer for 30 min at 37 °C. Tagmented nuclei were kept on ice before encapsulation.

#### scATAC-Seq library preparation and sequencing

Tagmented nuclei were loaded onto a ddSEQ Single-Cell Isolator (Bio-Rad). scATAC-Seq libraries were prepared using the SureCell ATAC-Seq Library Prep Kit (17004620, Bio-Rad) and SureCell ATAC-Seq Index Kit (12009360, Bio-Rad). Bead barcoding and sample indexing were performed with PCR amplification as follows: 37 °C for 30 min, 85 °C for 10 min, 72 °C for 5 min, 98 °C for 30 s, eight cycles of 98 °C for 10 s, 55 °C for 30 s and 72 °C for 60 s, and a single 72 °C extension for 5 min to finish. Emulsions were broken and products were cleaned up using Ampure XP beads. Barcoded amplicons were further amplified for 8 cycles. PCR products were purified using Ampure XP beads and quantified on an Agilent Bioanalyzer (G2939BA, Agilent) using the High-Sensitivity DNA kit (5067-4626, Agilent). Libraries were sequenced on HiSeq 2000 with 150 bp paired-end reads.

#### Sequencing reads preprocessing

Sequencing data were processed using the Bio-Rad ATAC-Seq Analysis Toolkit. This Toolkit is a streamlined computational pipeline, including tools for FASTQ debarcoding, read trimming, alignment, bead filtration, bead deconvolution, cell filtration and peak calling. The reference index was built upon the mouse genome mm10. For generation of the fragments file, which contain the start and end genomic coordinates of all aligned sequenced fragments, sorted bam files were further process with "bap-frag" module of BAP (https://github.com/caleblareau/bap). Downstream analysis was performed with SnapATAC [16], cisTopic[17] and ArchR[18] . Resulted sorted bam file of each sample was structured into hdf5 snapfiles (single-nucleus accessibility profiles) using snaptools version 1.4.1. Cell-by-bin matrices were created with bin size of 5,000bp. SnapATAC package was used to merge snapfiles based on the same bin coordinates. Peak coordinates obtained from Bio-Rad ATAC-Seq Analysis Toolkit were loaded on snapfiles to generate cell-by-peak matrix. Cell-by-peak matrix exported from SnapATAC was imported to create cisTopic object. Fragment files were used to create the Arrow file in the ArchR package.

#### Clustering and gene score/transcription factor activity analysis

We filtered out low-quality nuclei with stringent selection criteria, including read depth per cell (>2,000) and TSS enrichment score (>20%). Potential doubles were further removed based on the ArchR method. Bin regions were cleaned by eliminating bins overlapping with ENCODE Blacklist regions, mitochondrial DNA as well as the top 5% of invariant features (house-keeping gene promoters). Dimensionality reduction was performed using a nonlinear diffusion map algorithm available in the SnapATAC 1.0.0 package to produce 50 eigen-vectors of which the first 10 were selected in order to generate K Nearest Neighbor graph with K = 15. The clustering was performed using Leiden Algorithm available in the package leidenalg version 0.7.0 with a resolution of 0.4 and UMAP embedding were generated using umap-learn. ArchR was used to estimate gene expression for genes and TF motif activity from single cell chromatin accessibility data. Gene scores were calculated using the addGeneScoreMatrix() function with gene score models implemented in ArchR. addDeviationsMatrix() function was used to compute enrichment of TF activity on a per-cell basis across all motif annotations based on Chromvar.

#### cisTopic analysis

Topic modelling was performed using 10 to 35 (1 by 1), 40 and 500 iterations with cisTopics. The model was selected based on the highest log-likelihood. The cell-topic UMAP representation was obtained by using UMAP on the normalized topic-cell matrix (by Probability). Cell clustering result (6 clusters) from SnapATAC was used for visualization. Heatmap was then generated based on the cell-cisTopic distributions to identify topics associated with each cluster. To select a representative set of regions of the topic, Region-topic distributions were binarized with a probability threshold of 0.99 to identify the top contributing regions in each topic. For identifying enriched GO terms per topic, the binarized topics (i.e. sets of top regions per topic) was analyzed over GREAT or clusterProfiler[19].

##### Trajectory analysis

Trajectory analysis was performed in ArchR. addTrajectory() function in ArchR was used to construct trajectory on cisTopic UMAP embedding. To perform integrative analyses for identification of positive TF regulators by integration of gene scores with motif accessibility across pseudo-time, we used the correlateTrajectories() function which takes two SummarizedExperiment objects retrieved from the getTrajectories() function.

##### Enhancer analysis

To connect distal regulatory elements to promoters, we first used the addCoAccessibility() function in ArchR to calculate co-accessibility. To select links connecting distal regulatory element and promoter, two peaks in each pair were annotated by ChIPseeker separately and the links with only one peak falling in the promoter region were retained. To identify enhancers, datasets of H3K4me3, H3K27me3, H3K4me1, H3K4me2 and H3K27ac modifications in Id4-GFP-bright SSC were downloaded from GSE131656[20]. ChromHMM was used to train a 12 state model on all histone marks assayed. Chromatin states of DARs among different clusters, topic regions and distal element-promoter links were annotated using annotatr package [21], and candidate regions overlap "Active enhancer" states were considered as putative enhancers. ROSE was used for the identification of super-enhancers with H3K27ac dataset (GSE131656)[20]. Overlapping regions of the resulting sets of super-enhancers and CCANs were evaluated using makeVennDiagram in ChIPpeakAnno. CCANs and super-enhancers were associated with nearby genes using GREAT analysis with the default setting[22].

##### Footprinting analysis

Differential transcription factor footprints between control and UNC0638 treatment group were identified using the Regulatory Genomics Toolbox application HINT[23]. Aligned BAM files from control or UNC0638 treatment group were treated as pseudo-bulk ATAC-seq profiles and then subjected to rgt-hint analysis. Based on MACS calling peaks, we used HINT-ATAC to predict footprints with the “rgt-hint footprinting” command. We then identified all binding sites of a particular TF overlapping with footprints by using its motif from JASPAR with “rgt-motifanalysis matching” command. Differential motif occupancy was identified with “rgt-hint differential” command and ”–bc” was specified to use the bias-corrected signal.

#### Real-time PCR

Total RNA was extracted from cells by AllPrep DNA/RNA Micro Kit (QIAGEN). RNA was then converted to cDNA by reverse transcription using PrimeScript RT Master Mix (Takara). Real-time PCR was performed using Power SYBR® Green PCR Master Mix (Life Technology) following manufacturer's instructions. Primers used: Cdh1 forward (5'-3') CAGGTCTCCTCATGGCTTTGC, reverse (5'-3') CTTCCGAAAAGAAGGCTGTCC; Vim forward (5'-3') CCTTTACTGCAGTTTTTCAGG, reverse (5'-3') GTATTCTAGCACAAGATTTC.

#### Cell proliferation analysis

For each well of 24-well plate, $10^5$ cells were seeded and cultured in complete GSC medium. Each sample contains at least two technical replicates to minimize errors. The number of cells was counted on day 7 and day 14 of the experiment.

#### Cell invasion assay

Invasion assay was performed according to manufacturer's instructions (Corning). In brief, a layer of Matrigel (Corning) was coated on the transwell insert containing a polycarbonate membrane with 8 μm pores (Corning Costar). A single cell suspension containing 105 cells was seeded on the transwell membrane and the transwell insert was placed into a 24-well plate that contained complete medium. After incubation for 48 hours, cells in the upper chamber were removed and the transwell insert was fixed in 4% PFA for 30 minutes. Cells that migrated to the transwell membrane were stained with 0.1% crystal violet for 10 minutes and then observed under a light microscope. At least 6 fields were counted for each sample. The migration ratio was calculated by the area of cells divided by the total area of the visual field using the ImageJ software (NIH).

#### Spontaneous EMT assay and small molecule inhibitor treatment

Matrigel (Corning) was diluted with DMEM-F12 (Invitrogen) to reach a final concentration of 1-100 μg/ml. 250 μl of Matrigel solution was added to each well of a 24 well tissue culture plate and the plate was placed in 4°C overnight. The final Matrigel concentration was 0.1-10 μg/cm2. After coating overnight, the Matrigel solution was aspirated and the well was washed once with PBS before seeding cells. GSCs were seeded on Matrigel (0.1 μg/cm2) to induce domed-shaped colonies. We then transferred the colonies to Matrigel (10 μg/cm2) after which cells at the border could sense adjacent unoccupied space and spontaneously undergo EMT and migrate outwards. The proportion and distance that cells migrated were observed.

Small molecule inhibitors were immediately added after colony transfer and the cells were treated for 24 to 48 hours. RepSox (ab142139, Abcam), UNC0638 (ab142157, Abcam), Nintedanib (HY-50904, MedChemExpress LLC) and Erlotinib (SML2156, Sigma) were dissolved in DMSO. The final concentration of each inhibitor was: RepSox 5 μM, UNC0638 2 μM, Nintedanib 0.5 μM and Erlotinib 5 μM. Medium containing DMSO was used as a control.

#### In vitro small molecule inhibitor treatment and Fluorescence-activated cell sorting (FACS)

For small molecule inhibitor treatment subjected to flow cytometry, complete GSC media containing freshly diluted inhibitors were refreshed every 2 days. Medium containing DMSO was used as a control. Flow cytometry was performed after 4 days of drug treatment. For retinoic acid (RA) treatment, GSCs were cultured in complete GSC medium containing 0.5 μM RA (R2625, Sigma) for 20 hours before flow cytometry analysis. Medium containing DMSO was used as a control.

GSCs were subjected to flow cytometry analysis after trypsin digestion. For analysis, cells were incubated for 20 min at 4°C in staining buffer with Alexa Fluor 647 anti-mouse CDH1 antibody (147307, Biolegend) or PerCP-Vio700 anti-mouse CD90.2 antibody (130-102-204, Miltenyi Biotec). Flow cytometry was performed on a BD FACSAria Fusion Flow Cytometer (BD Biosciences). Data were analyzed by FlowJo software (FlowJo, LLC).

#### Immunofluorescence staining and confocal microscopy

Cells cultured on Matrigel-coated coverslips or testis sections on microscope slides were fixed in 4% (v/v) paraformaldehyde for 15 min at room temperature and rinsed three times in PBS for 5 min before staining. Cells were permeabilized by treating in PBS containing 0.2% Triton X-100 for 15 min. Cells were blocked for 2 hours in PBS containing 10% normal donkey serum (NDS) (Jackson Immunoresearch). Primary and secondary antibodies and respective dilutions were listed in Supplementary Table S7. Primary antibodies were applied overnight at 4°C. Appropriate secondary antibodies were applied for one hour at room temperature. Cell nuclei were counterstained with DAPI (1 μg/ml, Sigma). All confocal images were captured by Olympus FV1200 confocal microscope. FV10-ASW software (Olympus) was employed to create overlays of colors.

#### Statistical analysis

Statistical analysis. Assessment of statistical significance was performed using two-tailed unpaired t-tests, one-way ANOVA with Tukey multiple comparisons tests or Chi-squared tests. Statistical analysis was performed using GraphPad Prism v8. Associated P values are indicated as follows: *P < 0.05; **P< 0.01; ***P < 0.001; ****P < 0.0001; not significant (ns) P > 0.05.

### Supplementary Figures and Legends

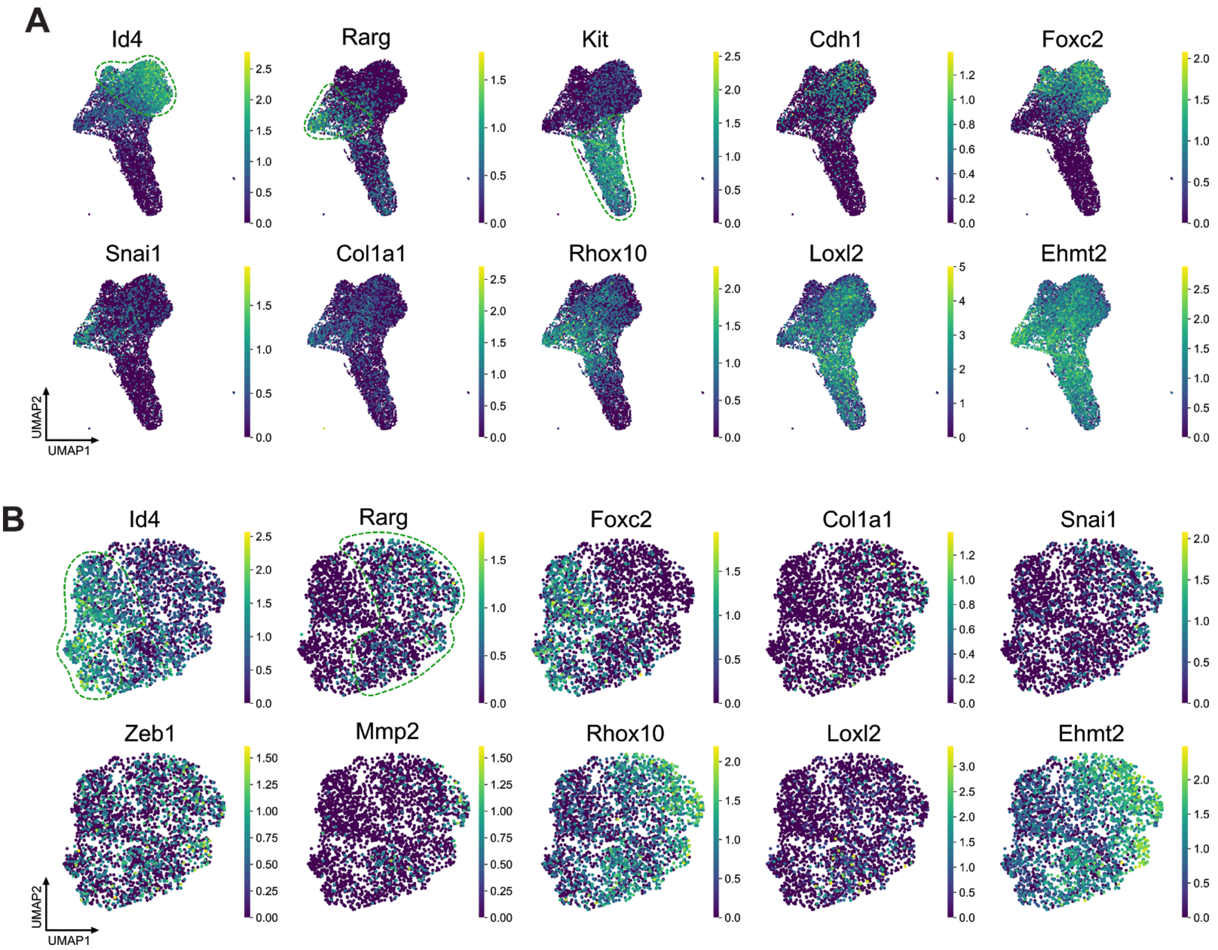


**Supplementary Figure S1. Gene expression heterogeneity in neonatal and adult undifferentiated spermatogonia revealed by scRNA-Seq, related to Figure 1.**
**A.** Gene expression level of representative ECM and EMT regulatory genes from scRNA-seq dataset of PND6 undifferentiated spermatogonia. Green dotted lines denote areas of the subpopulations for high *Id4* expression (stem cell-like), high *Rarg* expression (progenitor-like), high *Kit* (differentiating-like) respectively. **B.** Gene expression level of representative ECM and EMT regulatory genes from scRNA-seq dataset of adult undifferentiated spermatogonia. Green dotted lines denote areas of the subpopulations for high *Id4* expression (stem cell-like) and high *Rarg* expression (progenitor-like) respectively.

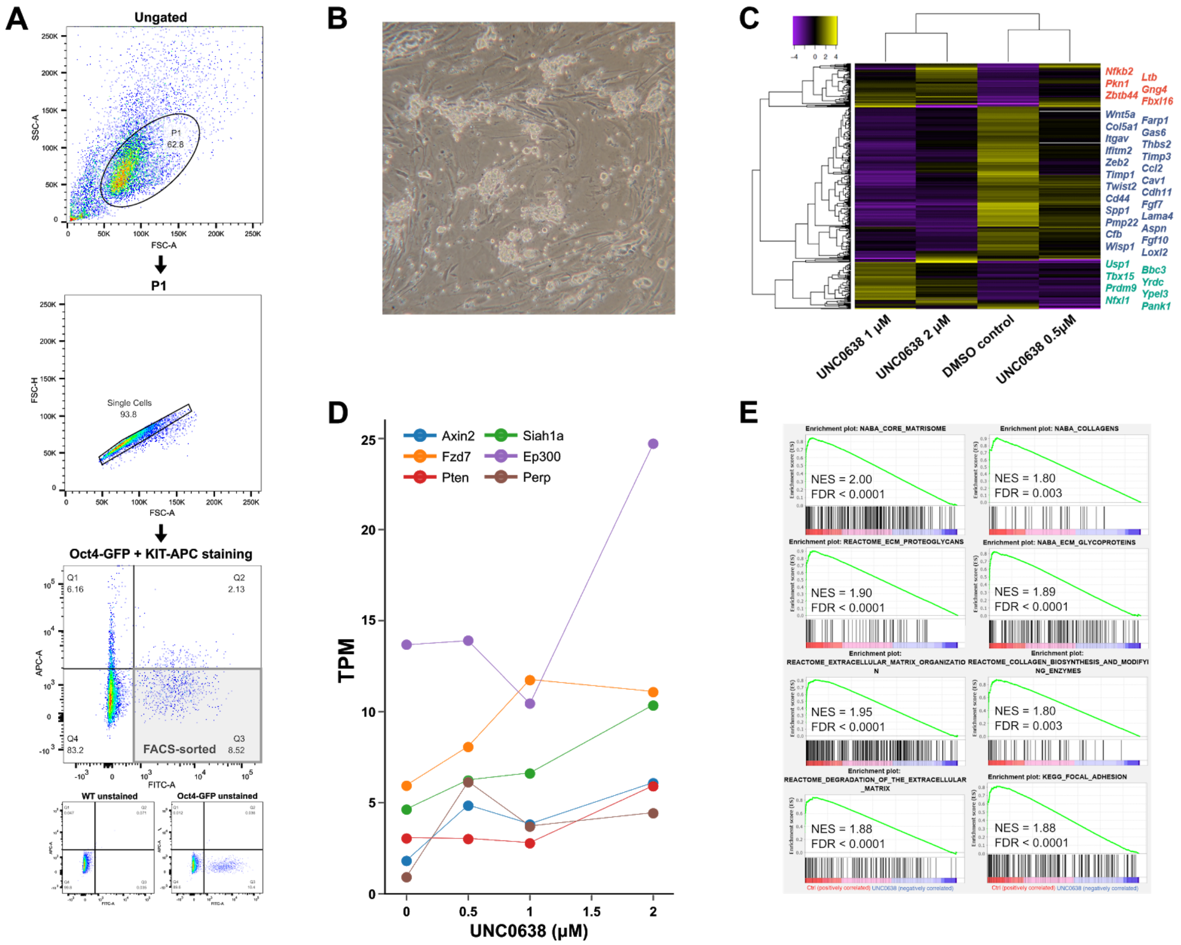


**Supplementary Figure S2. GSC derivation, culture, and gene expression characterization of G9a inhibition, related to Figure 1.**

**A.** Sorting strategy for isolation of undifferentiated (Oct4-GFP+/KIT−) SSCs at PND6. Oct4-EGFP positive cells were determined by comparing testicular cells from PND6 with wild-type control (WT). The KIT- population was gated using APC-KIT labelled sample compared with unstained control. Oct-GFP+/KIT- subpopulation (Q3) was FACS-sorted for culture. **B.** Bright-field images of GSC cultured on mouse embryonic fibroblast (MEF) layer. **C.** Heatmap of 435 differentially expressed genes identified by comparing UNC0638-treated groups and control groups ($p < 0.05$, FC > 2). **D.** Gene expression trend of selected upregulated genes upon different dosages of UNC0638 treatment. **E.** The top curated gene sets identified by GSEA showing enrichment in non-UNC0638-treated GSCs compared to UNC0638-treated GSCs were associated with ECM proteins and regulators.

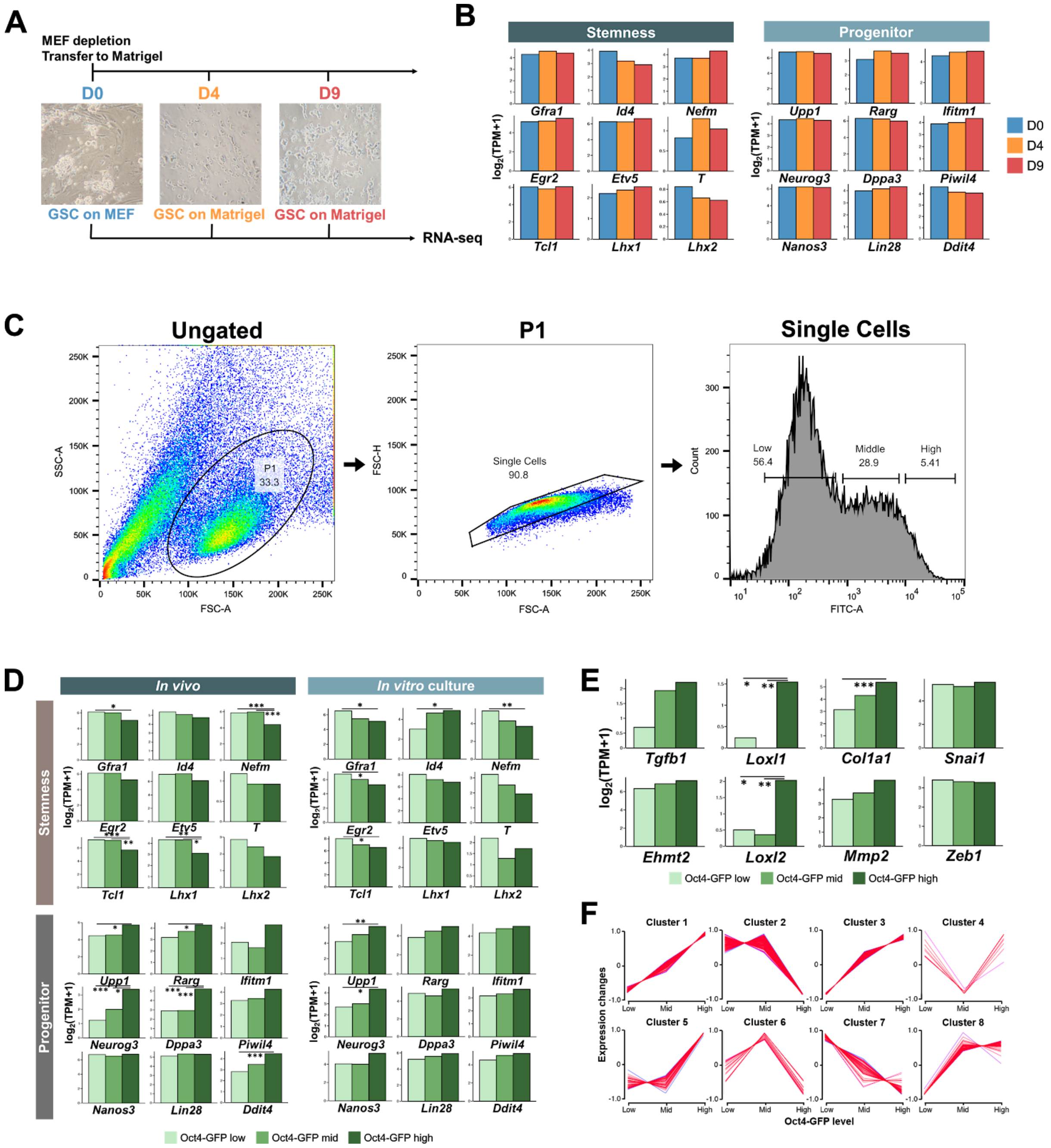


**Supplementary Figure S3. Gene expression characterization during MEF-to-Matrigel transfer and recapitalation , related to Figure 1.**

**A.** Schematic of GSC collection during ECM transfer from MEF to Matrigel. **B**. Bar plots showing the dynamics of stemness and progenitor gene expression in relation to MEF-to-Matrigel transfer. **C.** Sorting strategy for isolation of cultured Oct4-GFP-low/middle/high cells. **D.** Gene expression levels of representative stemness (top) and progenitor (bottom) genes in in vivo P6 SSC and in vitro P6 GSC culture. * FDR < 0.01. ** FDR < 0.05. *** FDR < 0.001. **E.** Gene expression level of representative ECM and EMT regulatory genes in in vivo P6 GSC and in vitro P6 GSC culture. * FDR < 0.01. ** FDR < 0.05. *** FDR < 0.001. **F.** Differentially expressed genes were identified among Oct4-GFP-low, Oct4-GFP-mid and Oct4-GFP-high groups and z-score-normalized ratios of the gene expression were subjected to soft clustering with Mfuzz software. The 8 clusters represent different expression profiles.

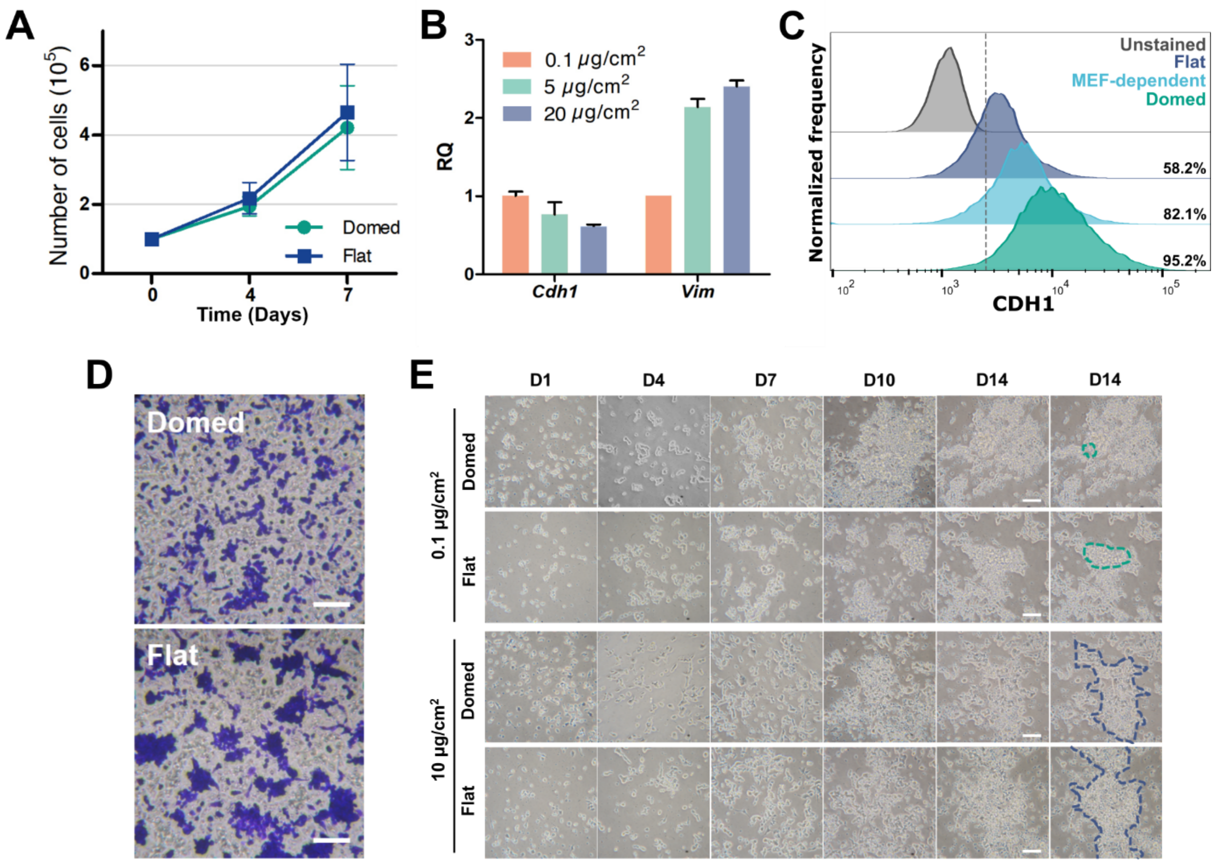


**Supplementary Figure S4. Characterization of the cellular property of domed and flat GSC colonies, related to Figure 2.**

**A.** Cell growth curve of domed and flat GSCs showed no significant difference on D7 and D14 respectively ($n \geq 3$). Significance was determined by an unpaired two-tailed t-test. **B.** RT-PCR analysis of expression of epithelial marker (Cdh1) and mesenchymal marker (Vim) of GSC cultured on different concentrations of Matrigel coating. Error bars are plotted with SD. **C.** Flow cytometry analysis of epithelial marker CDH1 in flat GSCs, MEF-dependent GSCs and domed GSCs. **D.** Matrigel-precoated membrane filter insert was used to measure in vitro invasiveness. After 48 hours of incubation, the domed or flat SSCs that migrated through the membrane were stained, and representative fields were photographed. Scale bars 200 µm. **E.** Bright-field images of GSC culture demonstrating the specific Matrigel concentration always leads to the same colony type, regardless of seeding domed or flat cells initially. Scale bars 100 µm.

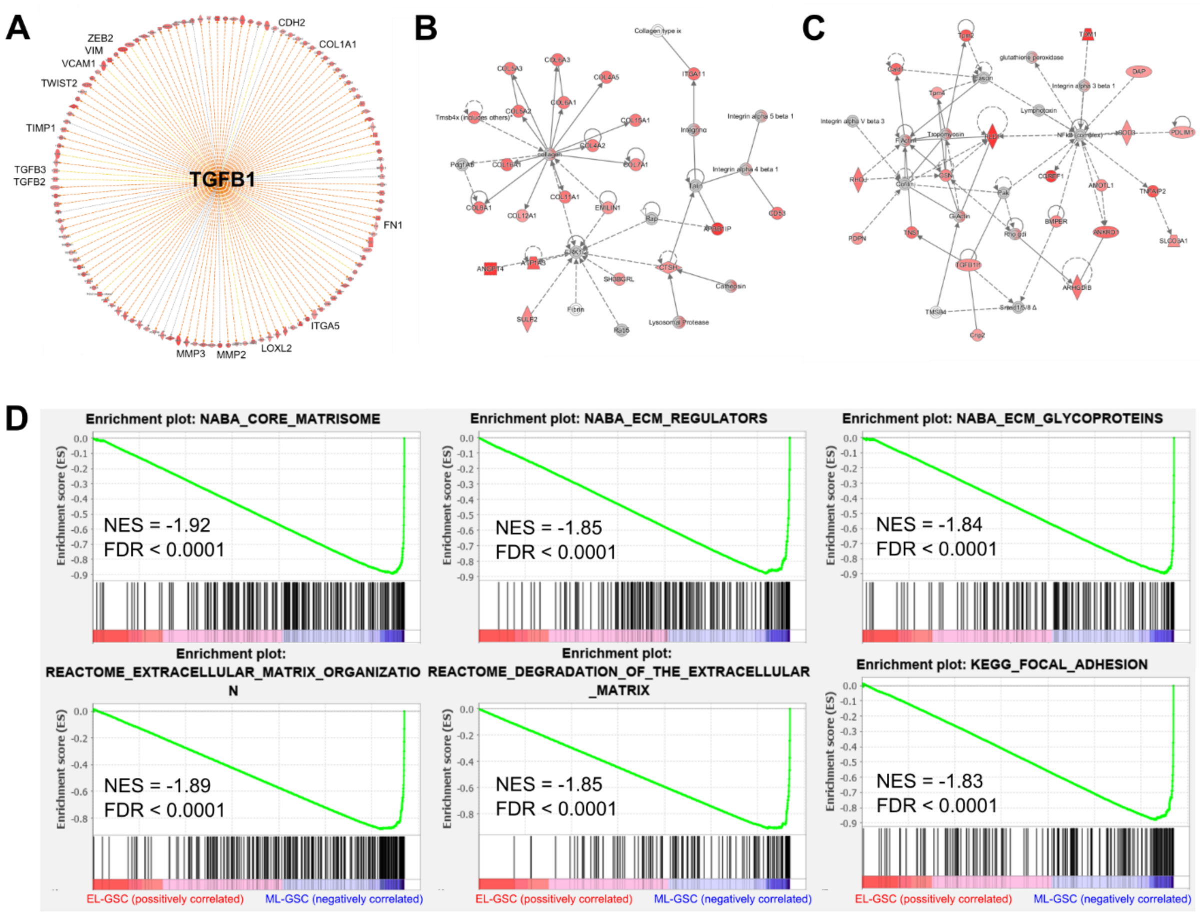


**Supplementary Figure S5. Ingenuity Pathway Analysis of epithelial-like and mesenchymal-like GSC, related to Figure 2.**

**A.** The network of the upstream regulator TGFB1 and its targeted genes (e.g. *Lox*, *Itga5*, *Mmp2*, *Vcam1*). Data illustrate TGFB1 as "activated" in ML-GSCs in upstream regulator analysis. Genes in red were greater expressed in ML-GSCs compared with EL-GSCs. An orange line indicates predicted upregulation, whereas the yellow line indicates expression being contradictory to the prediction. **B.** Topmost significant gene networks enriched in ML-GSCs. Associated network functions: Cancer, Connective Tissue Disorders, Organismal Injury and Abnormalities. Red indicates genes with increased measurement (Score: 35). **C.** Top second most significant gene networks enriched in ML-GSCs. Associated network functions: Cell Morphology, Connective Tissue Development and Function, Skeletal and Muscular System Development and Function. Red indicates genes with increased measurement (Score: 33). **D.** The top curated gene sets identified by GSEA showing enrichment in ML-GSC compared to EL-GSC were associated with ECM proteins and regulators.

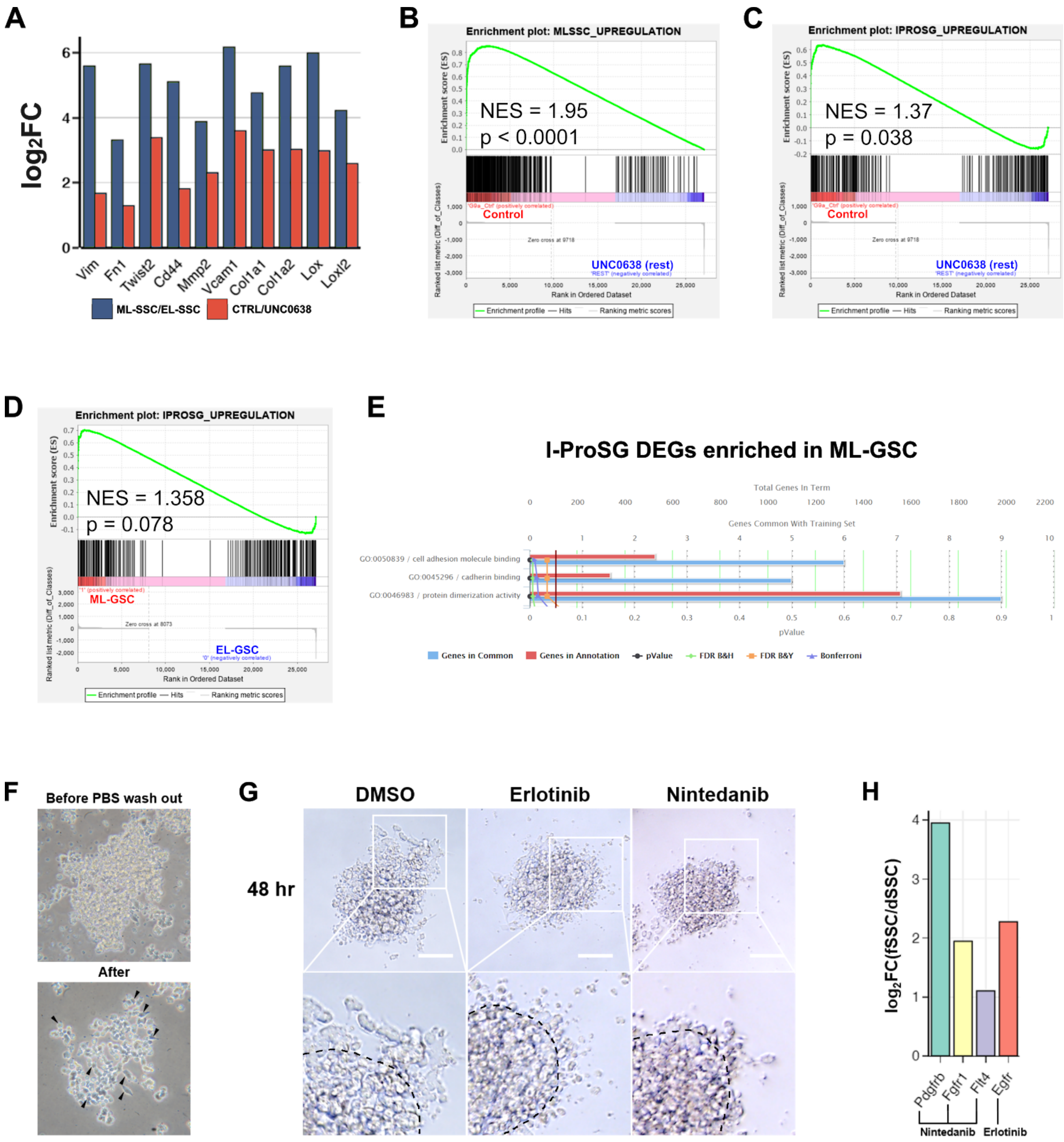


**Supplementary Figure S6. Comparison of mesenchymal-like GSC (ML-GSC), GSC after G9a inhibition and I-ProSG, related to Figure 2 and 3.**

**A.** Bar chart showing the expression change of representative overlapped EMT regulatory genes (upregulated in ML-GSC/downregulated after G9a inhibition). **B.** The gene set of ML-GSC upregulated DEGs showing enrichment in non-UNC0638-treated GSCs compared to UNC0638-treated GSCs. **C.** The gene set of I-ProSG DEGs (compared to other stages) showing enrichment in non-UNC0638-treated GSCs compared to UNC0638-treated GSCs. **D.** The gene set of I-ProSG DEGs to other germs cells showing enrichment in ML-GSC compared to EL-GSC. **E.** Gene Ontology (GO) terms significantly enriched in the enriched genes identified from **D**. **F.** In vitro spontaneous EMT assay. After seeding a domed-shaped colony on high-adhesion Matrigel, cells at the border could sense adjacent unoccupied space and spontaneously undergo EMT-like process (arrows). **G.** Bright-field images of domed GSC colonies treated with small molecule inhibitors (Erlotinib 5 μM, Nintedanib 0.5 μM) or vehicle (DMSO) for 48 hours in spontaneous EMT assay. **H.** Bar chart showing the expression change of genes encoding selected kinases upregulated in ML-GSC.

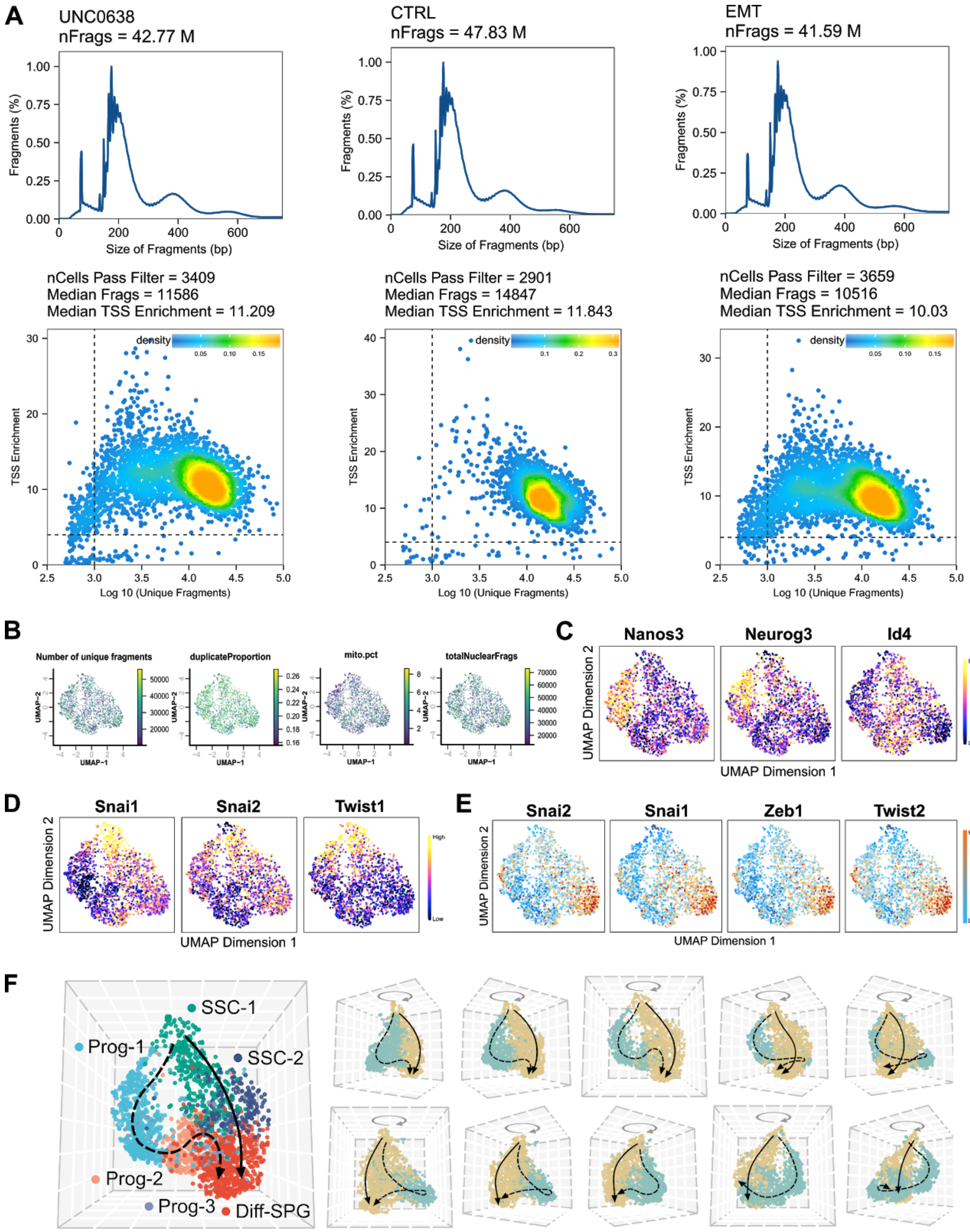


**Supplementary Figure S7. Single-cell chromatin state of GSC, related to Figure 4 and 5.** **A.** Quality assessment metrics for scATAC-Seq libraries. Fragment size distribution plot shows enrichment around 200 bp, indicating nucleosome-free and mono-nucleosome-bound fragments (top). Plot of TSS enrichment score versus the total number of unique fragments. Only cells lying in the upper right quadrant (marked by dashed lines) are retained for downstream analysis (bottom). **B.** UMAP projection of co-embedded cells colored by different QC metrics. **C.** Gene activity score of EMT marker genes shown in UMAP. **D.** Motif accessibility of EMT-TF motifs shown in UMAP. **E.** Gene activity score of SSC self-renew and differentiation genes are shown in UMAP. **F.** Identification of two trajectories correlated with cell states visualized by uniform manifold approximation and projection (UMAP) in 3D.

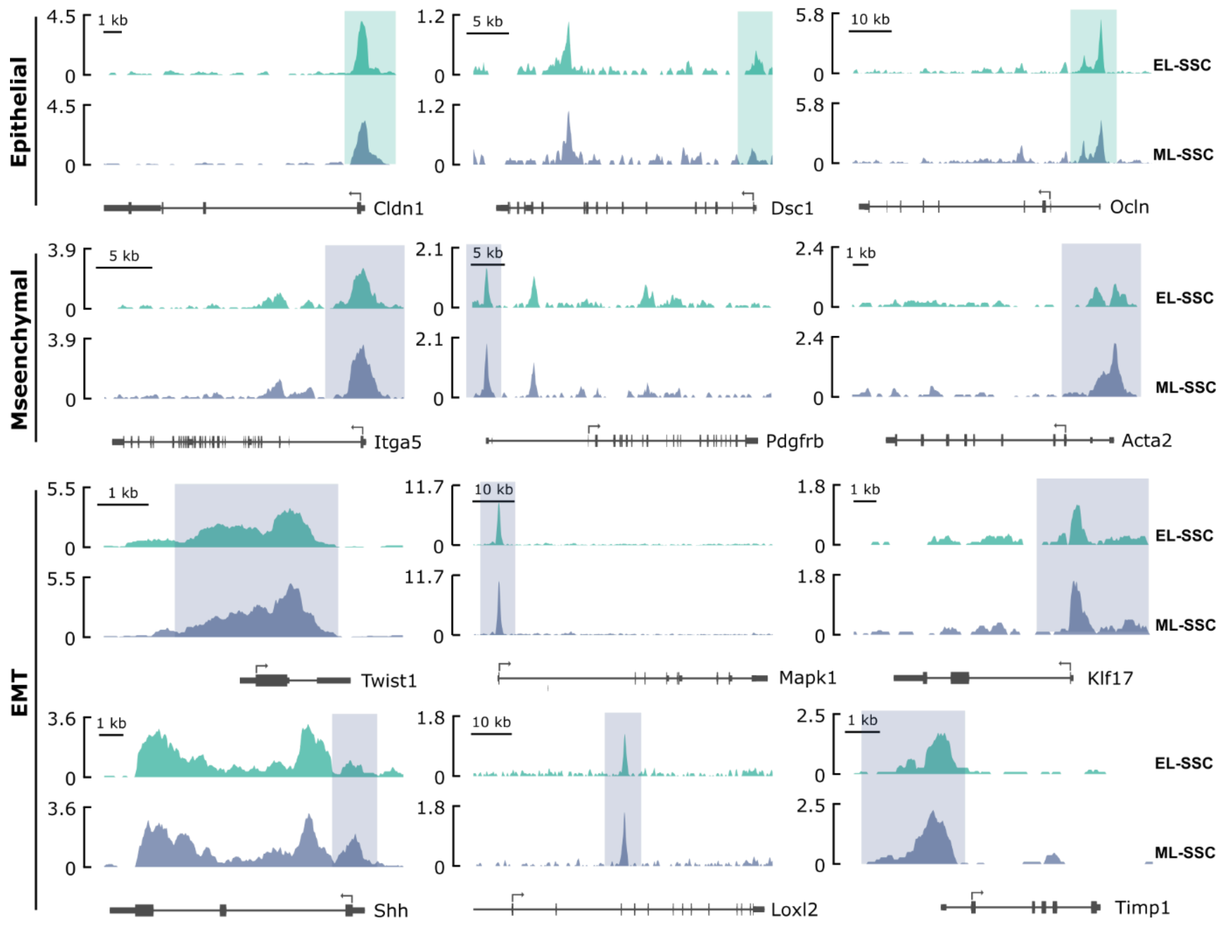


**Supplementary Figure S8. Bulk ATAC-seq of epithelial-like- (EL) and mesenchymal-like- (ML) GSC revealed chromatin accessibility of EMT-related genes, related to Figure 4.**

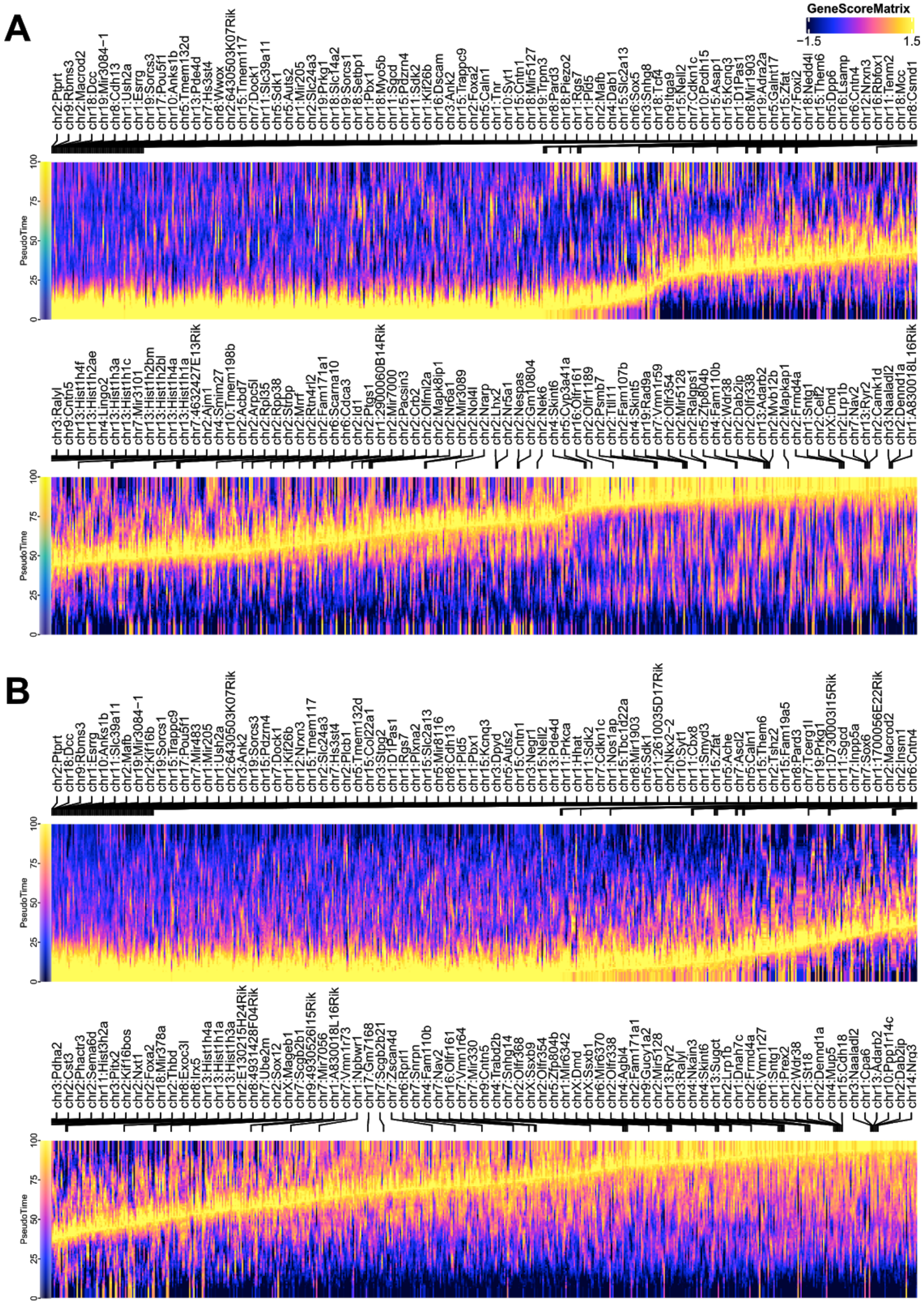


**Supplementary Figure S9. Genes with its gene score co-varies across pseudotime trajectory, related to Figure 5.**

Smoothened heatmap depicting TFs whose gene score co-varies across pseudotime of **A.** Trajectory 1 and **B.** Trajectory 2.

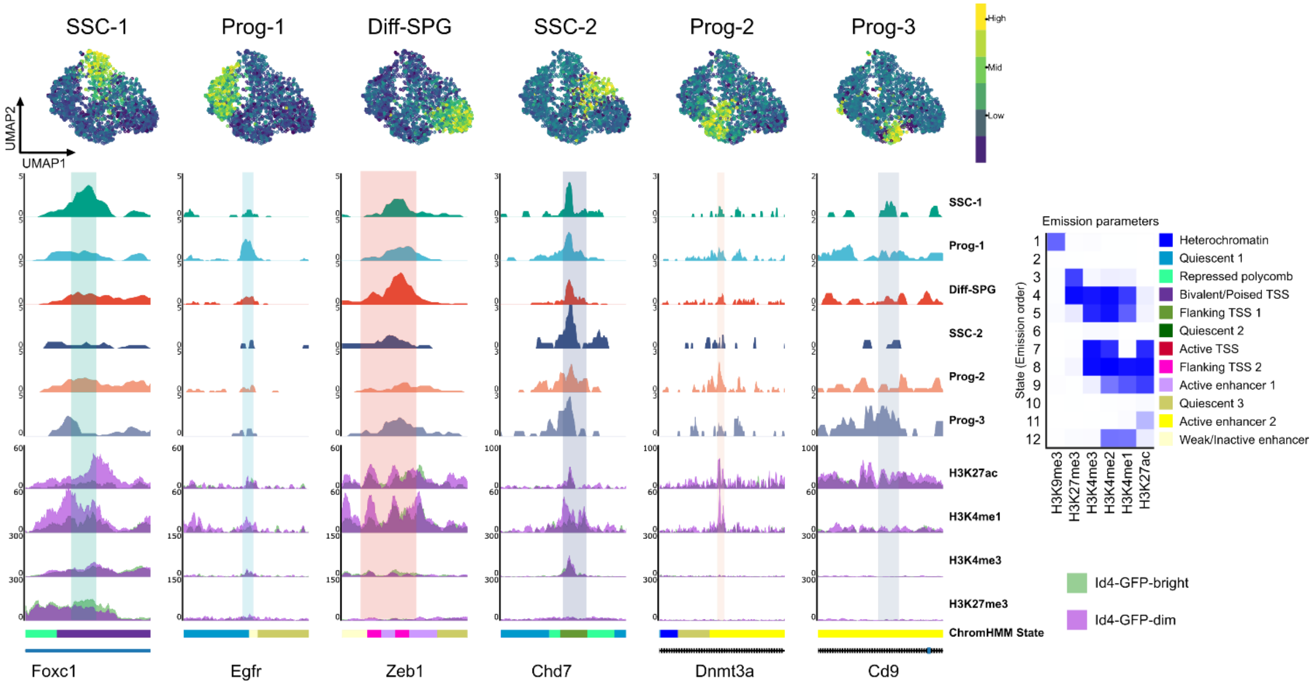


**Supplementary Figure S10. Cluster-specific enhancer regions of each spermatogonia subset, related to Figure 6.**
cisTopic cell UMAP color-coded by the normalized topic score of each spermatogonial subset (Upper panel) and respective aggregated scATAC-seq profiles showing one selected differentially accessible region for each subset (Lower panel). The selected gene is related to its function in spermatogonia.

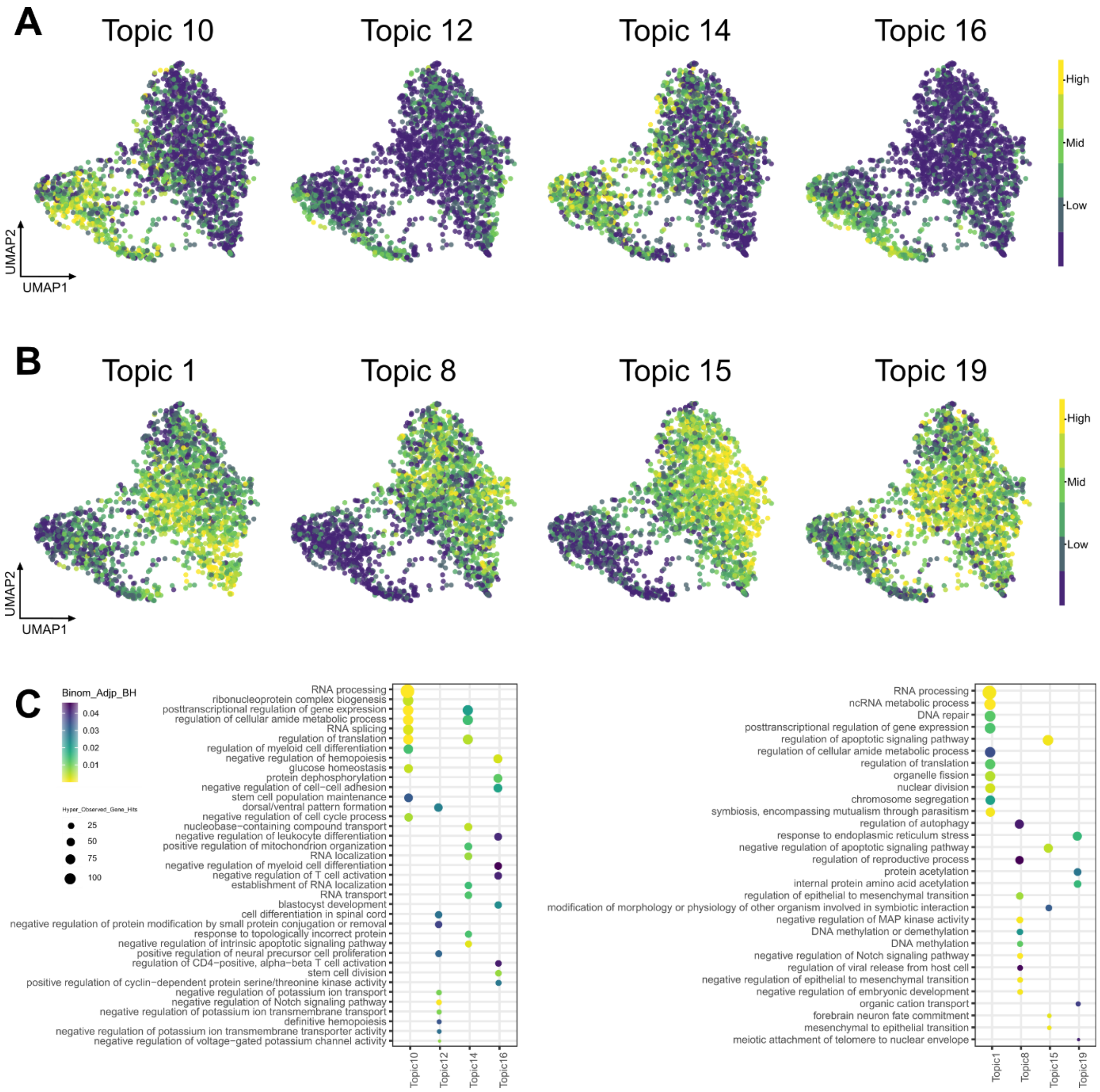


**Supplementary Figure S11. Regulatory topics related to Prog-2 and non-Prog-2 subsets, related to Figure 6.**

**A.** cisTopic cell UMAP color-coded by the normalized topic score of topics enriched in Prog-2 subset. **B.** cisTopic cell UMAP color-coded by the normalized topic score of topics enriched in non-Prog-2 subset. **C.** GREAT analysis of regions included in topics in **A** (Left panel) and **B** (Right panel).

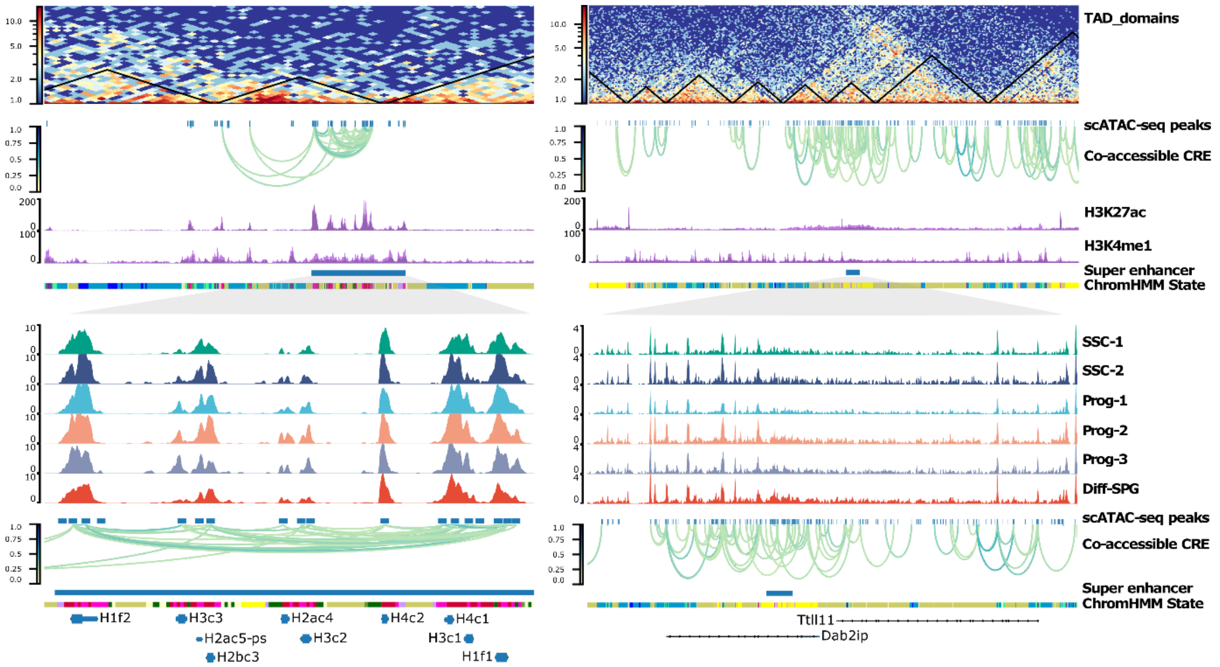


**Supplementary Figure S12. Super-enhancer (SE) regions, related to Figure 7.** Aggregated scATAC-seq profiles showing SE regions near *Hist* locus (Left panel) and *Dab2ip* (Right panel) with inferred TAD domains and co-accessible CREs.

### Supplementary Tables S1-S7

**Supplementary Table S1.** DEG list of GSCs treated with 0 μM (DMSO control), 0.5 μM, 1 μM and 2 μM UNC0638. Gene expression level was shown in $log_2$TPM.

**Supplementary Table S2.** Mfuzz clustering results of DEGs from Supplementary Table S1.

**Supplementary Table S3.** DEG list of Oct4-GFP-high, Oct4-GFP-middle and Oct4-GFP-low GSCs. Gene expression level was shown in $log_2$TPM.

**Supplementary Table S4.** Mfuzz clustering results of DEGs from Supplementary Table S3.

**Supplementary Table S5.** DEG list of epithelial-like (EL) and mesenchymal-like (ML) GSCs. Gene expression level was shown in $log_2$TPM.

**Supplementary Table S6.** GSC culture medium composition.

| | 10ml media | Work concentration | Storage | Company |
|---|---|---|---|---|
| StemPro | 8.55ml | - | 4C | Life Technologies |
| Human GDNF | 20ul | 40ng/ml | -80C | R&D Systems |
| Human bFGF | 10ul | 10ng/ml | -80C | BD |
| Mouse EGF | 5ul | 20ng/ml | -80C | Life Technologies |
| StemPro-Nutrient Supplement | 0.26ml | - | -80C | Life Technologies |
| BSA | 200ul | 0.20% | -20C | MP Biochemicals |
| GlutaMax | 100ul | 1x | -20C | Life Technologies |
| ES-FBS | 100ul | 1% | -20C | Life Technologies |
| Peni/Strep | 100ul | 1× | -20C | Life Technologies |
| NEAA | 100ul | 1× | -20C | Life Technologies |
| Sodium pyruvate | 100ul | 1× | -20C | Life Technologies |
| Transferrin | 100ul | 100μg/ml | -20C | Sigma |
| VITAMIN | 100ul | 1× | -20C | Sigma |
| D(+)-Glucose | 100ul | 1mg/ml | -20C | Sigma |
| D-biotin | 100ul | 10μg/ml | -20C | Sigma |
| Insulin | 25ul | 25μg/ml | -20C | Sigma |
| NaOH | 18ul | - | R.T. | Sigma |
| Progesterone | 15ul | 60ng/ml | -20C | Life Technologies |
| DL-Lactic acid | 10ul | 1μl/ml | -20C | Sigma |
| Ascorbic acid | 10ul | 100μM | -20C | Sigma |
| β-Estradiol | 10ul | 30ng/ml | -20C | Sigma |
| β-ME | 9.1ul | 50μM | 4C | Sigma |
| Putrescine | 6ul | 60μM | -20C | Sigma |
| Sodium selenite | 1ul | 30nM | -20C | Sigma |

**Supplementary Table S7.** List of antibodies and dilutions.

| | Company | Catalog # | Dilution | Application |
|---|---|---|---|---|
| **Primary antibody (clone)** | | | | |
| Rabbit anti-CDH1 mAb (24E10) | CST | 3195T | 1:200 | IF |
| Rat anti-vimentin mAb (280618) | R&D systems | MAB2105 | 1:50 | IF |
| Goat anti-uPAR pAb | R&D systems | AF534SP | 1:100 | IF |
| Rabbit anti-G9a mAb (C6H3) | CST | 3306T | 1:50 | IF |
| Rabbit anti-H3K9me2 pAb | Active Motif | 39041 | 1:250 | IF |
| Mouse anti-HES1 mAb (E-5) | Santa Cruz | sc-166410 | 1:1000 | Western blot |
| Mouse anti-GAPDH | Abcam | ab8245 | 1:1000 | Western blot |

| **Secondary antibody** | | | | |
|---|---|---|---|---|
| Rhodamine Red-X Donkey anti-Rabbit IgG (H+L) | Jackson Immunoresearch | 711-295-152 | 1:500 | IF |
| Alexa Fluor 647 Goat anti-Rabbit IgG (H+L) | Thermo Fisher | A-21244 | 1:500 | IF |
| Alexa Fluor 647 Goat anti-Rat IgG (H+L) | Thermo Fisher | A-21247 | 1:500 | IF |
| Alexa Fluor 647 Donkey Anti-Goat IgG (H+L) | Life Technologies | A21447 | 1:500 | IF |
| Mouse IgG HRP Linked Whole Ab | GE | NA931-100UL | 1:10000 | Western blot |
